# Quantifying Structural Diversity of CNG Trinucleotide Repeats Using Diagrammatic Algorithms

**DOI:** 10.1101/2020.05.30.124636

**Authors:** Ethan N. H. Phan, Chi H. Mak

## Abstract

Trinucleotide repeat expansion disorders (TREDs) exhibit complex mechanisms of pathogenesis, some of which have been attributed to RNA transcripts of overexpanded CNG repeats, resulting in possibly a gain-of-function. In this paper, we aim to probe the structures of these expanded transcript by analyzing the structural diversity of their conformational ensembles. We used graphs to catalog the structures of an NG-(CNG)_16_-CN and NG-(CNG)_50_-CN oligomer and grouped them into sub-ensembles based on their characters and calculated the structural diversity and thermodynamic stability for these ensembles using a previously described graph factorization scheme. Our findings show that the generally assumed structure for CNG repeats—a series of canonical helices connected by two-way junctions and capped with a hairpin loop—may not be the most thermodynamically favorable, and the ensembles are characterized by largely open and less structured conformations. Furthermore, a length-dependence is observed for the behavior of the ensembles’ diversity as higher-order diagrams are included, suggesting that further studies of CNG repeats are needed at the length scale of TREDs onset to properly understand their structural diversity and how this might relate to their functions.

**STATEMENT OF SIGNIFICANCE:** Trinucleotide repeats are DNA satellites that are prone to mutations in the human genome. A family of diverse disorders are associated with an overexpansion of CNG repeats occurring in noncoding regions, and the RNA transcripts of the expanded regions have been implicated as the origin of toxicity. Our understanding of the structures of these expanded RNA transcripts is based on sequences that have limited lengths compared to the scale of the expanded transcripts found in patients. In this paper, we introduce a theoretical method aimed at analyzing the structure and conformational diversity of CNG repeats, which has the potential of overcoming the current length limitations in the studies of trinucleotide repeat sequences.

## INTRODUCTION

Trinucleotide repeats and the disorders associated with their expansion is a growing field of study. Understanding the pathogenesis of trinucleotide repeat expansion disorders (TREDs) requires insights into the structure-function relationship that relates these repeat sequences to their RNA transcripts and the symptoms attributed to them. Trinucleotide repeats are microsatellites known to exhibit large length variability (1–3). Based on their locations within the gene and the extents of expansion, these repeats are known to cause a variety of neurological disorders such as Huntington’s disease and fragile X syndrome. It is often unclear whether the RNA transcript of the repeats themselves or another aberrant gene product is responsible for their cytotoxicity (4–7). Amongst the diverse family of TREDs, those caused by CNG repeats have become a major research target.

While expanded CNG repeats may lead to aberrant protein products with extended sequences of glutamines (polyQ), many of the trinucleotide expansions are not found in coding regions. In these cases, cytotoxicity may be due to the mRNA transcripts, either via a gain or loss of function. Often, disease manifestation is associated with a critical expansion threshold, and as a result, onset of disease appears to be a function of age. Examples of gain of function has been demonstrated in myotonic dystrophy type 1 (DM1) (4), where expanded CTG repeats in the 3’-untranslated regions of the *dystrophia myotonica* protein kinase gene produces a RNA transcript which interacts with CUG-binding proteins (CUGBP1) and muscleblind-like (MBNL1) proteins. These interactions alter protein levels in the cell, which in turn affects their function as splicing regulators, leading to symptoms (8–10). Understanding the *in vivo* structure of the repeats may lead to a better understanding of how RNA with expanded CNG sequences may interact with these proteins.

The structures most often associated with the gain of function hypothesis for CNG expanded RNA sequences cited in the literature is a necklace-like structure composed of a long stretch of successive two-way junctions interposed by shorts helixes and with a hairpin stem-loop cap (11–16). Many of the studies conducted, however, are based on CNG repeat oligomers, whereas the threshold of TRED disease onset is typically associated with expansions of 60 to 100 units or longer. Additionally, the structures resolved are limited to those which can be isolated and crystalized. As the length of the CNG repeats grow, the diversity of accessible structures could grow rapidly as well, with the necklace structure comprising only one possible subset of motifs out of many. This leaves a gap in our understanding of the structure-function relationship that may be responsible for pathogenesis in CNG-related TREDs. The structural diversity of CNG chains on the order of lengths comparable to the critical expansion thresholds of TREDs remain unclear. In this paper, we present a calculation aimed at addressing this question using an algorithm based on a diagrammatic approach.

## MATERIAL AND METHODS

### Backbone Conformational Entropy and Secondary Structures

An open RNA strand is characterized by an ensemble of many diverse conformations and is in a high-entropy state. If this RNA sequence can fold and develop secondary structure(s), the entropy of the chain will decrease as the chain folds because the number of conformations that are consistent with the secondary structures in the folded state is necessarily lower than the unfolded state. This decrease in conformational entropy *S* of a RNA can be viewed as a result of the constraint(s) imposed by the secondary structural elements present in the fold on the possible conformations of the chain. The entropy loss from the unfolded state to the folded state is:

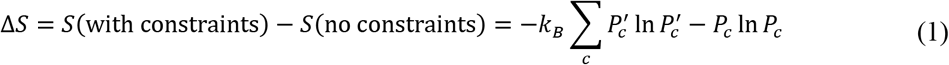

where *c* is a chain conformation, *P_c_* and 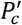 are the normalized probabilities of that conformation with and without the constraint(s) imposed by the secondary structure, and *k_B_* is Boltzmann’s constant. If an alternative fold with a different set of secondary structural elements exists, it will in general have a different conformational entropy because the constraints are different. Adding more constraints to the chain in the form of more complex secondary structures will necessarily lead to a more negative Δ*S*, but the constraints imposed by the different secondary structural elements in a fold are in general not independent, e.g. Δ*S* with constraints A + B is not necessarily equal to Δ*S* with constraint A plus Δ*S* with constraint B.

In order to more easily characterize RNA secondary structures, Schlick et al. have proposed a diagrammatic scheme (17–19). Fig. 1 shows examples of some of the diagrams representing different kinds of secondary structural elements. The bottom row of Fig. 1 (a) illustrates the diagram of a three-way junction. Each of the three helices is represented by a black dot. Unpaired loops are represented by curved lines. The bottom row of Fig. 1(b) shows the diagrammatic representation of a pseudoknot. The two black dots now represent the two paired regions, whereas the unpaired loops are represented by straight or curved lines. A triplex is illustrated in Fig. 1 (c). A black triangle is used in its diagrammatic representation to represent the three-base interaction in this secondary structural element. Fig. 1(d) shows a quadruplex, and a black square is used to represent the interactions of the four bases in this secondary structure. In these diagrams, the points where two or more lines converge are called “vertices”. The lines emanating from each vertex are called “edges” and their number define the degree *d_υ_* of vertex *υ*. A dot (representing a duplex) always has four edges and *d_υ_* = 4. A triangle (representing a triplex) always has six edges with *d_υ_* = 6, and a square (representing a quadruplex) always has eight edges and *d_υ_* = 8. Because all RNAs are linear polymers, any graph representing a RNA fold is necessarily Eulerian (20), meaning there is a way to trace through the entire diagram over all its edges only once. This also implies that either the degree of every vertex will be even or only two vertices will be odd while all other are even. In such Eulerian graphs, the number of edges *E* including the two dangling ends on the 5’ and 3’ ends is

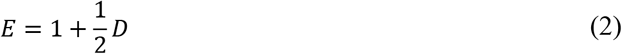

where *D* = Σ_*υ*_*d_υ_* is the total degree over all vertices in the diagram.

**Figure 1.**
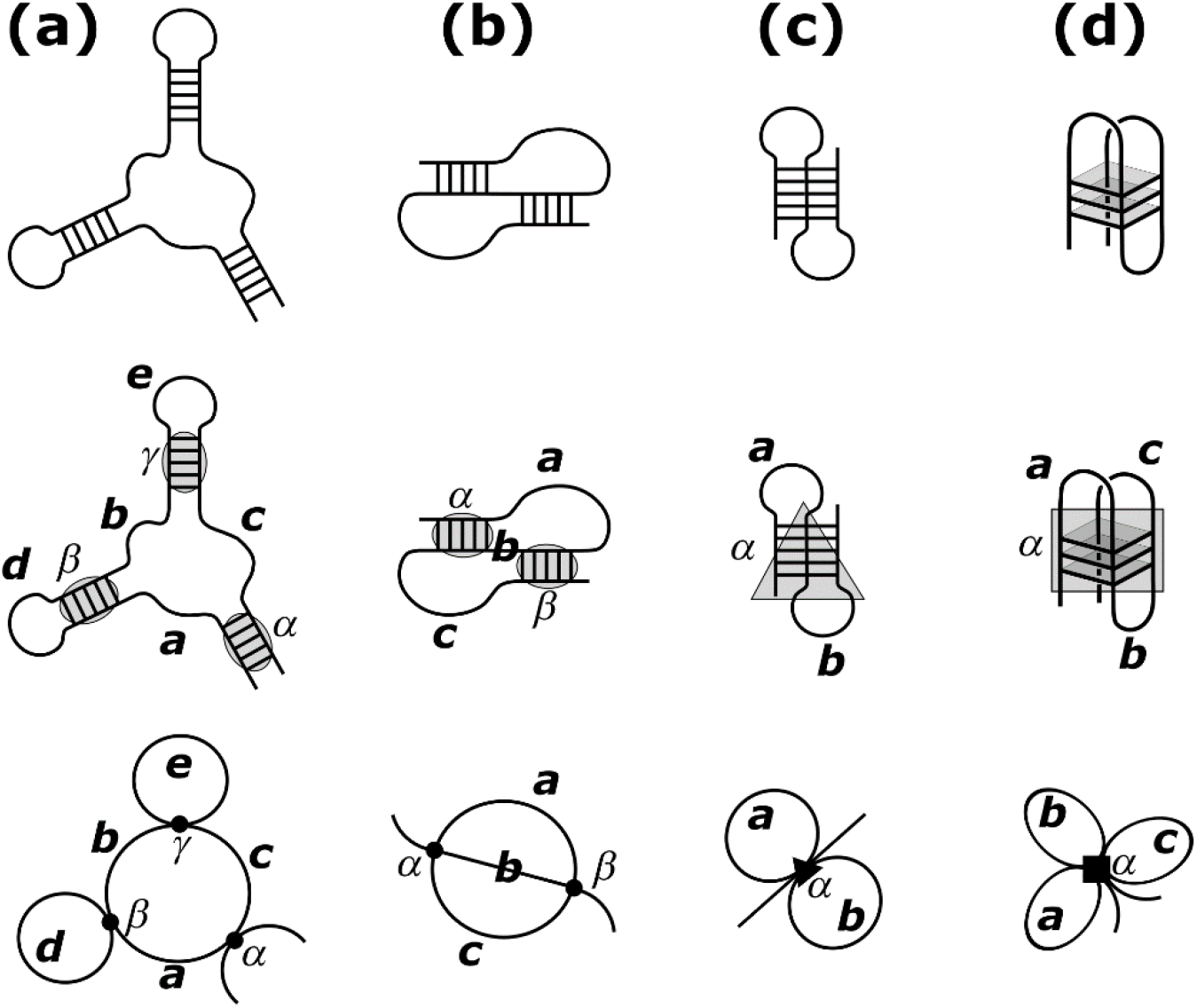
Examples of diagrams representing different secondary structural elements. (a) A three-way junction where each dot represents a helix and loops are represented by lines. (b) A pseudoknot where black dots represent paired regions and unpaired loops are represented by lines. (c) A triplex structure represented diagrammatically by a black triangle. (d) A quadruplex represented by a black square. Figure adapted from (21).

In addition to their utility in characterizing RNA secondary structures, the diagrammatic representation proposed by Schick et al. is also useful for the calculation of the conformational entropy of folded RNA structures. In a recent paper (21), we described how the constraints imposed by the secondary structural elements of any folded state can be broken into approximately independent sets using a factorization strategy based on how the elements of the diagram are connected. An example of how this factorization works is illustrated in Fig. 2 for a three-way junction. A “fragile vertex” is defined as any vertex that if removed from the diagram disconnects it into two or more disjoint pieces. Fig. 2 shows that all three vertices in the diagram of a three-way junction are fragile. Disconnecting the diagram at these fragile vertices generates the factorized diagrams on the far right of Fig. 2. When a diagram is completely factorized, it breaks up into irreducible pieces. We have proven that the conformational entropy of the fold is also approximately separable when a diagram is reducible, and Δ*S* becomes the simple sum of the entropies of all the irreducible pieces. For example, the total entropy Δ*S* of the three-way junction in Fig. 2 can be reduced to the sum of the three closed diagrams on the right. The big circle with arcs labeled *a, b* and *c* represents the unpaired loops in the three-way junction. The two smaller circles labeled *d* and *e* represent the loops in the hairpins. The edges corresponding to the 5’ and 3’ ends of the folded structure contribute nothing to Δ*S* since they contain no additional constraints and can be omitted from the completely factorized diagram shown on the right. The dots represent duplexes of different helix length (*α, β*, or *γ*). Each fragile vertex contains additional enthalpic and entropic free energy contributions depending on its size. The free energy of each fragile vertex can, for example, be estimated using Turner’s nearest-neighbor model (22–24), data from computer simulations, or other experimental data.

**Figure 2.**
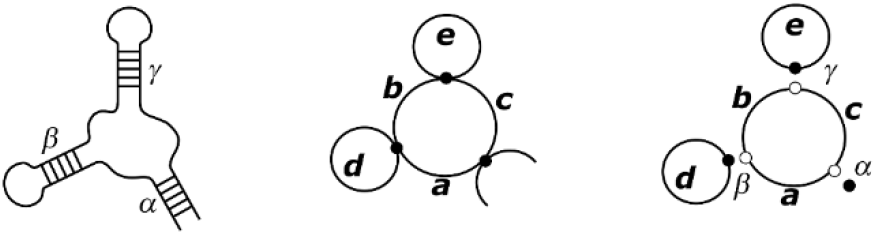
Factorization of the constraints in a three-way junction into approximately independent contributions. The total entropy Δ*S* is reduced to the sum of the three closed diagrams on the right.

### Monte Carlo Simulations

Diagrammatic factorization provides a simple recipe for the calculation of the conformational entropy of any RNA fold. To make use of it, a library of conformational entropy data must be compiled for every irreducible element representing various types of secondary structural elements (hairpin, junction, duplex, triplex, quadruplex, etc.) of different sizes, as well as for other unfactorizable structures such as pseudoknots. This library can be sourced from experimental data or from computer simulations. In a previous paper (21), we have provided a complete and consistent set of Monte Carlo simulation results for the entropy values of hairpins, two-, three- and four-way junctions. These correspond to diagrams in which the vertices are all degree 4 (i.e. dots). While some of the same data are available from melting experiments (22, 24, 25), not everything is. We have relied on extensive computer simulations to compile an internally consistent data set.

This paper further extends this data library, adding results for non-canonical base pairs and diagrams for quadruplexes and pseudoknots. These new data also serve to demonstrate factorizability questions in pseudoknots and quadruplexes, highlighting comparisons and contrasts with what has already been proven for two-, three- and four-way junctions. This data library provides entropic contributions to the free energy associated with the unpaired loops of each diagram. Free energies of formation of the paired regions associated with the vertices in each diagram are independent of the edges in these MC simulations, and they are added back in separately as needed. Vertex free energies are not included in the reported data for hairpin initiation, pseudoknot formation, and quadruplex formation obtained from direct simulation.

Monte Carlo (MC) simulations have been carried out using our in-house Nucleic MC program for high-throughput conformational sampling of RNAs (26). Detailed discussions of the mixed numerical/analytical treatment and closure algorithm used in simulating the sugar-phosphate backbone(26–30) and accounting for steric interactions (31,32), solvent effect (32–34), and counterions’ influence (35, 36) have been presented in previous publications. Using Nucleic MC, we generated thermal ensembles consisting of several million uncorrelated conformations for chains with many different secondary-structural constraints corresponding to a number of different classes of diagrams.

To evaluate the conformational entropy of quadruplexes and pseudoknots, poly-U constructs of many different structures were simulated. For diagrams involving helices with Watson-Crick (WC) base pairs, long hairpin structures in the protein databank were melted to obtain the appropriate starting conformation. Using the same parameters for defining base pairing events from our previous study (21), we then identified and counted spontaneous base pair formations during the simulation to measure the entropic cost of initiating any new base pair constraint within the structure. Multiple base-pairing constraints are associated with some of these structures. To evaluate the entropy of these, we computed the entropy cost for forming the first constraint, and holding the first constraint, we then computed the additional entropy cost for forming the second constraint, etc. Since entropy is a state function, any thermodynamic pathway between the initial (open) and final (folded) states will yield the correct Δ*S*. For example, the starting structure of a pseudoknot was chosen to produce the proper length for the *α* helix as shown in Figure 1, with the size of the seeded hairpin chosen to provide a range of lengths in the final assembled pseudoknot structure.

For quadruplexes, parameters for identifying Hoogsteen base pair are needed. The structures of pyrimidine-purine base pairs utilizing the purine’s Hoogsteen edge (PDBID 1GQU (37), 2QS6 (38), 1K2G (39), and 2H49 (40)) as well as quadruplexes with different topologies (PDBID 1KF1 (41) and 143D (42)) were used to define the base pairing criteria for identifying Hoogsteen pairs. These selection parameters for WC and Hoogsteen pairs are summarized in Fig. 3.

**Figure 3.**
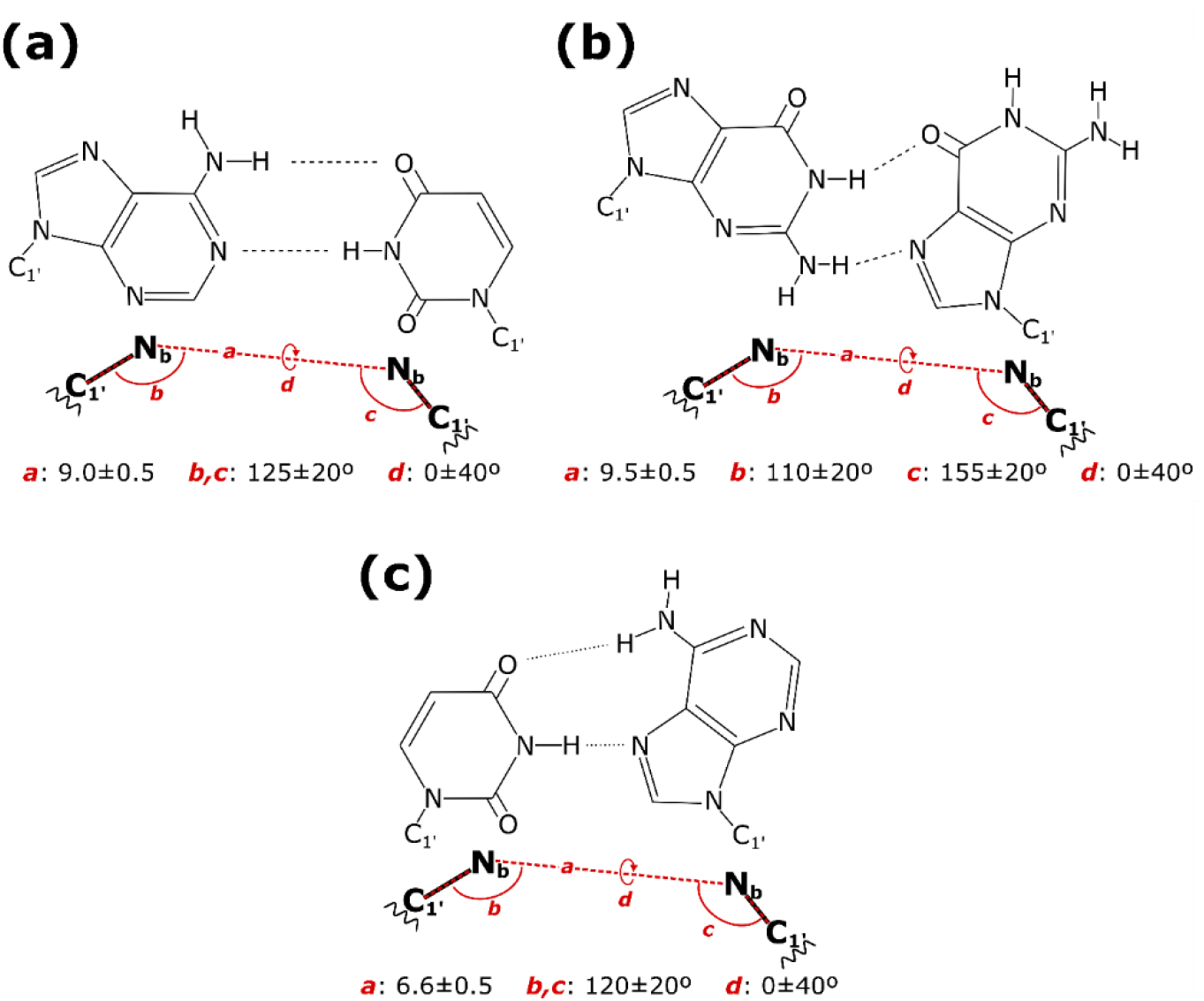
Geometric criteria used in defining (a) Watson-Crick base pairing geometry (43), (b) G-G Hoogsteen geometry (41, 42), and (c) purine-pyrimidine Hoogsteen geometry (41, 44).

### Evaluating Conformational Ensembles of CNG Repeats

Using the factorization scheme described above and a library of the entropy values of the irreducible elements, the entropy of the conformational ensemble of any RNA sequence can be evaluated. We illustrate this using the CNG repeat sequence 5’-NG(CNG)_8_CN-3’ as an example. Figs. 4(a) through (d) show four possible secondary structures of this sequence, and their corresponding diagrammatic representations are shown next to each. If we consider only WC base pairs, the longest uninterrupted canonically paired duplex length in CNG repeat sequences is only 2 base pairs (bp). These are highlighted by the blue boxes in Fig. 4. In the corresponding graphs, these are represented by blue dots. Since the structure in (a) has four 2-bp duplexes whereas (b) only has three, the total vertex free energy of (a) should be lower because base pairs are stabilizing. But on the other hand, (b) has fewer secondary structural constraints than (a), and therefore (b) is expected to have a more favorable conformational entropy. In an equilibrium ensemble of this sequence, we expect a thermodynamic competition between maximizing the number of vertices versus maximizing the diversity of the conformational ensemble. Furthermore, as our data library disallows hairpins shorter than 3-nt, the structure in (a) is the only conformation consistent with the diagram shown in (a). However, for structure (b), there are multiple alternative structures consistent with the graph shown in (b). These alternative structures can be obtained by permuting the junctions among the various positions along the sequence. For example, permuting the two junctions in the asymmetric bulge leads to a different structure without affecting its topology. Also, transposing a subsegment within one junction with another junction produces a different structure without altering the topology. For example, one can remove a single (CNG) unit from the hairpin and transpose it into the first junction, making both 4-nt long, to derive a new structure with a symmetric bulge instead of the asymmetric one in (b) without altering the topology. The entropy associated with the configurational diversity of the topological class represented by the graph in Fig. 4(b) is therefore higher than (a) and favors (b) over (a). In addition to (a) and (b), there are many other structures for the same sequence which belong to other topological classes. Fig. 4(c) and (d) show two additional examples. The structure in (c) corresponds to a three-way junction, while the structure in (d) consists of a pseudoknot plus a hairpin. Whereas (a) and (b) have different number of 2-bp duplex units, structures (b), (c) and (d) all have three duplexes. Because of this, structures (b), (c) and (d) have approximately the same vertex free energy and their competition for relevance within the conformational ensemble of this sequence is controlled by entropy alone. The goal of this study is to quantify the size and diversity of these CNG repeat ensembles using the diagrammatic techniques described above. Notice that each structure is characterized by a certain number of nucleotide (nt) units *ℓ* distributed over the unpaired regions among the loops and junctions, which in the graphs are associated with *E* edges. We will see that the problem of calculating the entropy is equivalent to finding all the possible ways of distributing the *ℓ* unpaired nucleotides over the *E* edges in the diagram.

**Figure 4.**
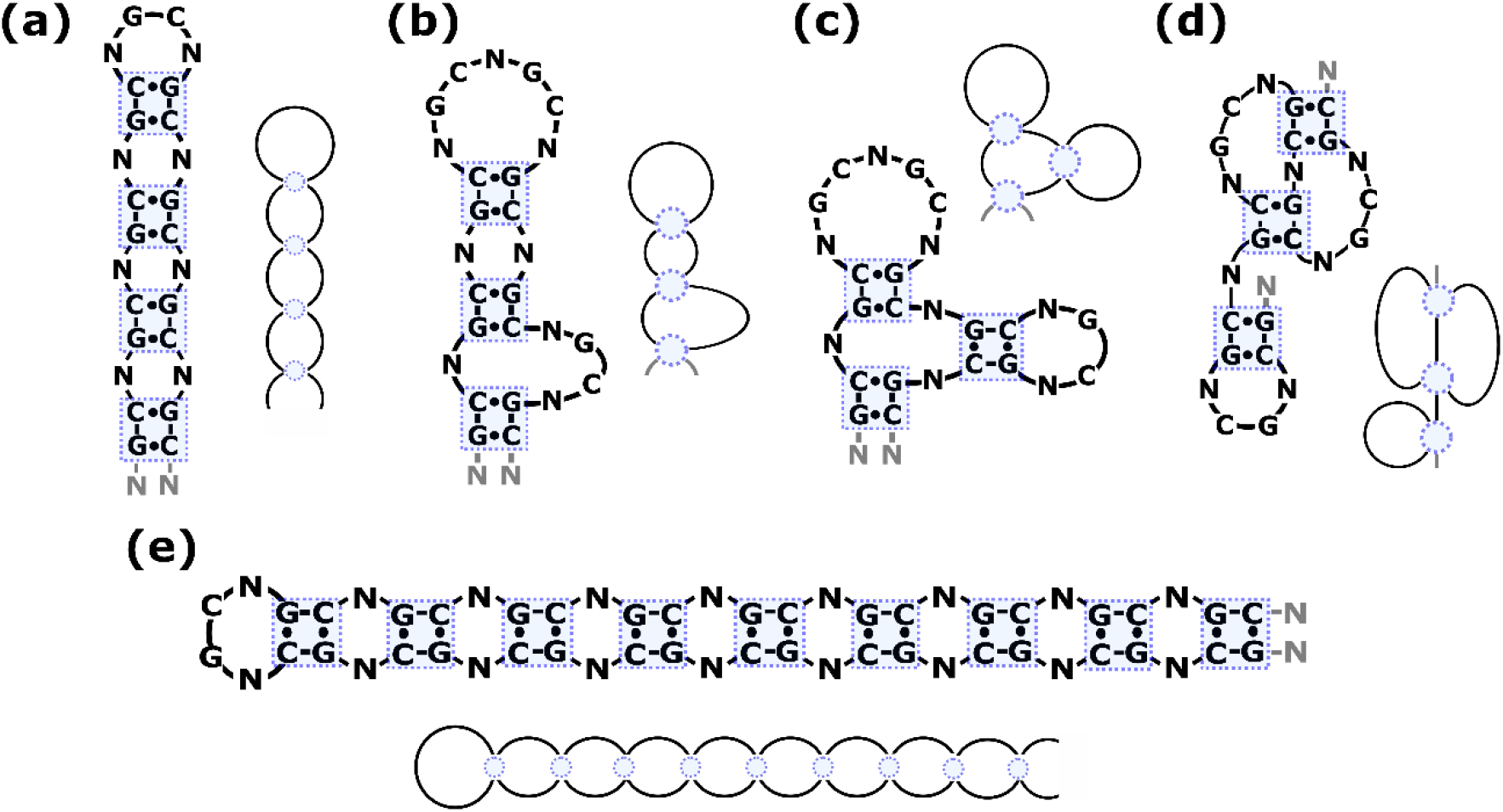
(a-d) Examples of different structures of the 5’-NG(CNG)_8_CN-3’ repeat sequence belonging to distinct topological classes. (e) Maximal hairpin structure of 5’-NG(CNG)_19_CN-3’.

To evaluate the volume of the conformational ensemble of a 5’-NG(CNG)_M_CN-3’ repeat sequence of a certain length *M*, we divide the ensemble into subsets according to the total degree of the graphs. Recall that the total degree *D* of a graph is equal to the sum of the degrees over all its vertices. Using this definition, the total degree of the graph in Fig. 4(a) is 16 because each vertex corresponding a duplex is degree 4, since 4 edges emanate from it. On the other hand, the graphs in (b), (c) and (d) all have total degree *D* = 12, because of the three duplexes present in each of those structures. Earlier, we have also mentioned that the total degree of a graph is related to the number of edges *E* in it by *E* = 1 + *D*/2. Because of this, we now recognize that even though the graphs in Fig. 4(b), (c) and (d) all belong to distinct topological classes, they all have the same number of edges because they have the same total degree. The diagrams in Fig. 4(b), (c) and (d) all have 7 edges because they are all degree 12. Furthermore, since the vertices are fixed-length duplexes, the total lengths of all the edges for all diagrams of degree *D* are also the same for sequences containing the same number of (CNG) repeats *M*. For example, the structures in Fig. 4(b), (c) and (d) all have total edge lengths of *ℓ* = 16 nt.

In a previous paragraph, we described how the entropy of a certain class of diagrams is derived from the permutation of the edges and the transposition of subsegment lengths among the edges. Restating this more precisely in terms of combinatorics, the structural diversity of a certain topological class is related to the number of possible ways in which the total edge length in a structure consisting of ℓ nts can be distributed among the *E* edges in the diagram. Because of this, grouping diagrams by total degree is advantageous compared to grouping them according to topological class. Since diagrams of the same degree also have the same number of edges, the combinatoric problem is identical for diagrams across the same degree, regardless of which topological class they belong to. This allows us to recycle the solution of the same combinatorics problem on diagrams of many different classes, as long as their total degrees are the same. This also means that the intrinsic diversities of the different subsets of the ensemble represented by different topological classes of graphs belonging to the same total degree are identical. The only difference between two topological classes belonging to the same total degree lies in the conformational entropies of the irreducible elements, which are different for different types of secondary structures. For example, the probability of observing an 8-nt hairpin loop in the ensemble of all possible conformations is very different from that of observing an 8-nt loop inside a three-way junction, even though they are both loops of the same length. These different entropy values are supplied by the library we complied using the MC simulations described above.

We summarize our solution to the combinatorics problem with a few useful equations here. Referring to Fig. 4, notice that each edge segment has a minimum length of 1 nt and can vary only by multiples of 3 nt. Therefore, the length of the *i*-th edge in a diagram can be represented by *1* + 3*j_i_*, where *j* is a non-negative integer, and there are *E* of these. The chain, which takes the form 5’-NG(CNG)_M_CN-3’, contains *M* + 1 (CNG) repeats. We always assume both the 5’ and 3’ ends have a dangling N nucleotide, so the total length of interest for the sequence is (3*M* + 4) nt. A degree-*D* diagram has *D* nts in the duplexes because all base pairs come in stacks of 2, so the total edge length is *ℓ* = (3*M* + 4) – *D*. The number of edges is 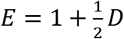. In terms of this, *ℓ* = 3(*M* + 2 – *E*) + *E*. Subtracting the minimum length of 1 nt for each of the *E* edges, the number of transposable nts is 3(*M* + 2 – *E*), but they must occur in 3-nt multiples. Therefore, the combinatorics problem is reduced to finding all sets of non-negative integers {*j*_1_,*j*_2_,⋯*j_E_*} such that 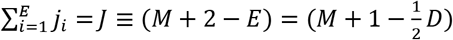, which also implies that the maximum theoretical degree for a chain with *M*+1 repeats is 2(*M* + 1). The process of dividing up the nucleotides into the edges of the graph is equivalent to creating a string of *E* non-negative integers with zeroes allowed such that they sum to *J*. This is the problem of determining all weak compositions of *J* in combinatorics; for a graph which has *E* edges, this is the enumeration of all weak *E*-composition of *J* (20, 45). For the purposes of this study, the enumeration is done by brute force with the correct compositions being stored as a valid structure of the graph ensemble. The collection of valid structures *α* for each graph Ξ forms an ensemble with each structure contributing a weight, 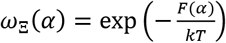, corresponding to its inherent free energy cost *F*(*α*) to the partition function of the graph ensemble, *Z*(Ξ) = Σ*ω*_Ξ_(*α*). For each graph ensemble, we can then define the ensemble-averaged conformational cost, 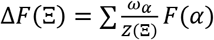, and the entropy 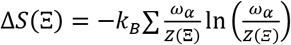. All graph ensemble calculations were carried out for two different RNA strands, NG-(CNG)_16_-CN and NG-(CNG)_50_-CN. For brevity, these will be referred to by their full length *n*, which are 17 and 51 respectively, in the result section.

## RESULTS

### Loop Initiation Entropies Involving Hoogsteen Pairs

Previously, we reported data for the entropies of initiating hairpin loops of different sizes seeded by WC pairs. To initiate a *n*-nt loop, the constraint associated with the base pair suppresses the conformational diversity of the backbone, which suffers an entropy penalty leading to a free energy cost Δ*G*(*n*) which increases with the loop length *n*. The free energy of formation of loops utilizing WC base pairing geometry are shown in Fig. 5 as the black filled circles. RNA triplexes and quadruplexes, on the other hand, must use noncanonical base pairing geometry on their Hoogsteen edges to initiate loops. The geometric constraints on the backbone needed to facilitate a purine-purine Hoogsteen pair (e.g. G:G) or a purine-pyrimidine Hoogsteen pair (e.g. A:C) are shown in Fig. 4(b) and (c), respectively. These Hoogsteen-specific geometric constraints produce higher free energy requirements for loop initiation compared to loops formed via WC interactions. Fig. 5 shows MC results for initiation free energies needed to form a loop using G:G Hoogsteen geometry (solid blue squares) and loops formed via purine:pyrimidine (R:Y) Hoogsteen geometry (open squares). Both types of Hoogsteen-pair geometry loops require higher free energy compared to WC-pair initiated hairpins. Note that these values pertain only to the entropic cost placed on the chain to close the loop and additional contribution from the base pairs themselves are added separately as needed.

**Figure 5.**
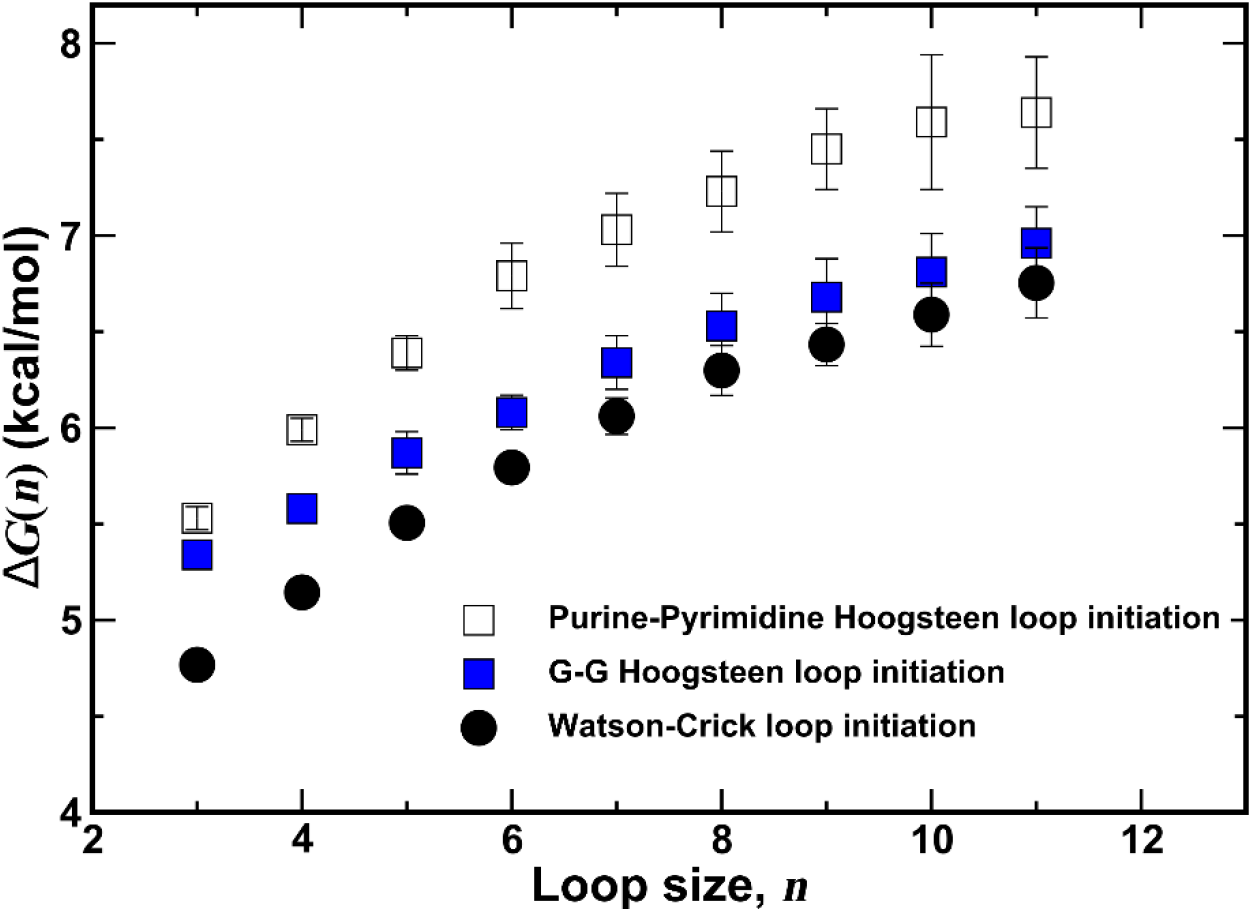
Comparison of loop initiation using different sets of base pairing criteria. In comparison to the Watson-Crick initiation cost, the purine-pyrimidine Hoogsteen initiation costs are effectively shifted up by a constant. This is consistent with the difference in the targeted inter-base distance and the almost identical range of bond and torsion angles. The G-G Hoogsteen loop initiation, despite sharing the same inter-base distance, suffers a larger cost at small loop lengths that is associated with the base positioning and the different inter-base angles.

### Quadruplexes

Quadruplex structures on DNA have been observed on d(GGG-NNN)_n_ repeats (46). These G-quadruplexes typically consist of a triple-deck sandwich of four Gs on each layer, interacting with each other via G:G Hoogsteen pairs. The d(NNN) sequences act as linkers, connecting the vertices of the triple sandwich. Various linker topologies have been identified. These are exemplified by the structures found in PDB IDs 1KF1 (41) and 143D (42). In 1KF1, the linkers are threaded through the G-quadruplex structure connecting the bottom corner of one edge of the triple sandwich with the top of an adjacent edge. In 143D, the linkers are threaded by connecting either the bottom corner of one edge with the bottom corner of an adjacent edge, or the top corner with the top corner of an adjacent edge.

The type of quadruplexes that are most relevant to 5’-NG(CNG)_M_CN-3’ RNA repeats are the double-deck sandwich structures illustrated by Fig. 6. Instead of three layers, the quadruplex structure in Fig. 6 has only two layers. The linker topology shown in Fig. 6 follows a bottom to top threading pattern, analogous to 1KF1. (CXG)_n_ repeats where X=G can potentially produce quadruplex structures of the type shown in Fig. 6, with each linker being either 1-nt (-C-), 4-nt (-CGGC), 7-nt (-CGGCGGC-) in length, or even longer.

**Figure 6.**
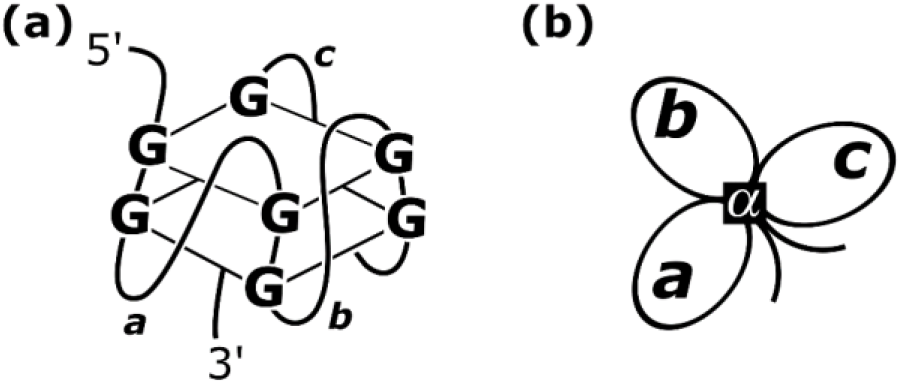
(a) A possible quadruplex structure relevant to CNG repeat sequences. The structure’s entropy is determined by the three loops labeled *a*, *b* and *c*. (b) Diagrammatic representation of a quadruplex, showing its dependence on the three loop lengths *a*, *b* and *c*, as well as the number of layers *α*.

There are three linker loops in a quadruplex structure. These are labeled *a*, *b* and *c* in Fig. 6 in the 5’ to 3’ direction. The entropic free energy costs for initiating the first loop *a*, the second loop *b* and the third loop *c* to connect the G on the bottom of one edge of the quadruplex to the G on the top of the next edge are tabulated in Table 1 for a double-deck quadruplex structure. In the MC simulations, loop *a* was initiated first. After this loop was formed, the free energy of initiating loop *b* was computed by holding the first two edges of the quadruplex fixed. After loop *b* was formed, the free energy of initiating loop *c* was then computed by holding the first three edges of the quadruplex fixed. The results in Table 1 suggest that as the linked loops get longer, the free energy cost of forming the loop also increases. This trend is not dissimilar to that observed in Fig. 5 for the hairpin initiation free energies. But as the quadruplex structure was assembled, the loop free energies also become progressively higher from *a* to *b* to *c*. This is presumably due to increased steric congestion in the core of the quadruplex structure, making it more difficult for loop *b* to form compared to *a*, and in turn more difficult for loop *c* to form compared to *b*. For linker loop *c*, the frequency of observing its formation in the MC simulations were too rare to be able to determine their free energies accurately for lengths > 5 or < 2 nt, and these have been left out of Table 1. In addition to the loops *a*, *b* and *c*, there are entropic penalties associated with constraining the backbone to the four edges of the quadruplex. The total free energy cost for this is also given in Table 1.

**Table 1.**
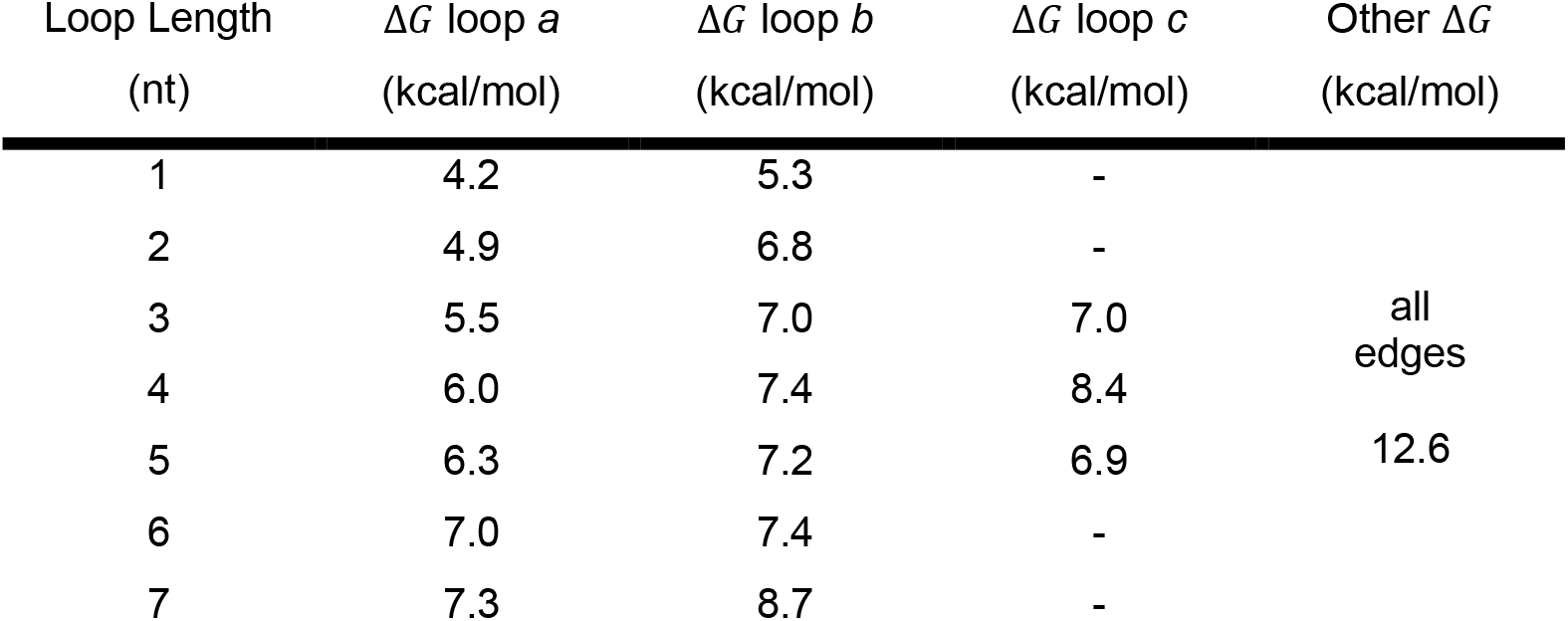
Cost of Initiating the Loops of Fig. 6. Initiation free energies for the *a*, *b* and *c* loops inside a double-deck quadruplex structure from MC simulations. The typical statistical error on each value is approximately ± 0.05 kcal/mol.

Previously, we found that vertices in graphs such as the helix in a hairpin or a stem in a 2-, 3-, or 4-way junction divide diagrams into approximately independent pieces. For the quadruplex, this independence is not strictly obeyed, because as Table 1 shows, the initiation free energies of the *a*, *b* and *c* loops are asymmetric with respect to exchange and they are no longer independent of each other. To accommodate this, we can modify the graph factorization scheme by simply redefining the entire quadruplex structure together with its *a*, *b*, and *c* loops as one irreducible element, instead of assuming the loops are separable. Since quadruplexes typically have very limited loop lengths, this does not affect the validity or impact the utility of the graph factorization scheme described above.

### Pseudoknots

The conformational entropy of a pseudoknot is determined by the lengths of the three loops a, b, and c, as well as the duplex lengths α and β. In the pseudoknot structures most relevant to CNG repeat sequences, α and β are 2 bp, the interhelix length b is 1 nt, and the loop lengths a and c are 1,4, 7,…. An example of how such pseudoknots fit into a (CGN) repeat sequence is shown in Fig. 4(d). Prior studies in the literature by Cao and Chen suggest that the three loops of a pseudoknot can be treated independently (47–49) when considering their entropies. Our simulation results for the pseudoknot structures most relevant to CNG repeats corroborate this.

To calculate the conformational entropy costs for pseudoknots, we calculated the cost for each steps in a thermodynamic pathway that folds a free chain into the final pseudoknot structure, passing through a hairpin structure along the way (an example of such pathways for the a=b=c=1 case can be found in Fig. S1 in the Supplemental Information). Consequently, the formation of the pseudoknot’s three loops in our method is due to a single base pairing event which turns an existing hairpin structure into a pseudoknot, and this can happen on either the 5’ side or the 3’ side. Additionally, there are extra entropy costs for extending the helices to reach their target lengths α and β. In Fig. 7, we summarize the entropic cost and standard error of forming the entire pseudoknot for the four smallest pseudoknot structures relevant to (CNG)_n_ repeats. The cost is calculated as the sum of costs to go from an open chain to the appropriate hairpin and then from the hairpin to the final pseudoknot structure relative to the cost of two 2-bp duplexes. Multiple pathways connecting the initial open chain and the final structure were used and the averages are shown in Fig. 7. The map of the pathway used for each of the structure can be found in the Supplemental Information as Fig. S1-S4, and the costs for each step of the pathways can be found in the Supplemental Information as Table S1-S4.

**Figure 7.**
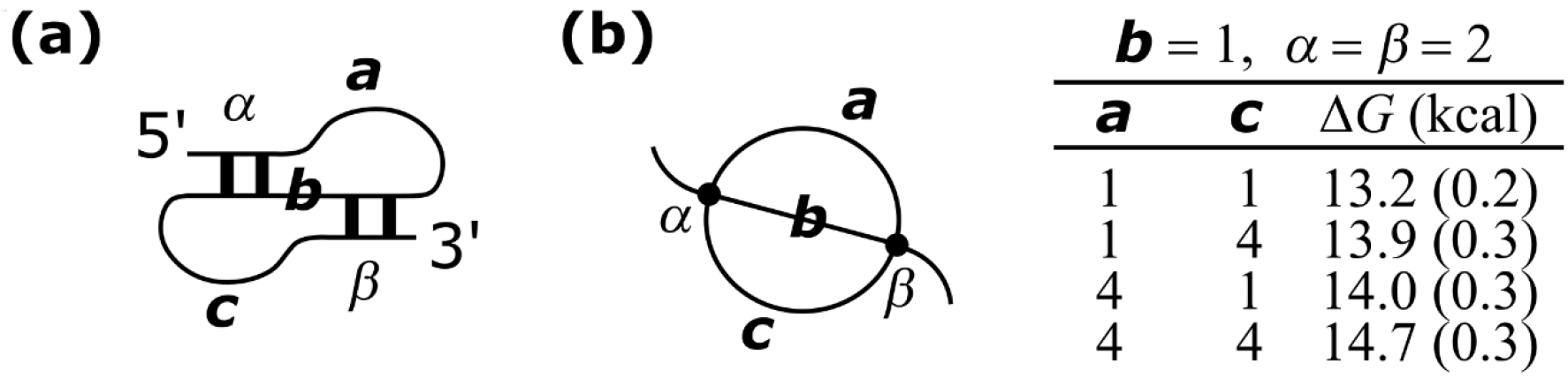
(a) The conformational entropy of a pseudoknot is determined by the lengths of the three loops labeled *a*, *b* and *c*, as well as the duplex lengths *α* and *β*. In the pseudoknot structures most relevant to CNG repeat sequences, *α* and *β* are both 2 nt, the interhelix length *b* is 1 nt, and the loop lengths *a* and *c* are 1,4, 7,… (b) Diagrammatic representation of a pseudoknot and calculated average cost for the four smallest pseudoknot structures relevant to CNG repeats.

### Ensembles of CNG Repeats

Given a 5’-NG(CNG)_M_CN-3’ repeat sequence, we first partitioned the conformational space according to the total degree of the diagrams, and then by collecting all accessible folded structures which share the same graph representation into a subset ensemble. For each structure within a subset ensemble, we calculated its weight using the entropic costs of all its irreducible elements as determined by the library derived from our MC data. The ensemble’s partition function of the subset represented by a graph Ξ was then used to calculate its sub-ensemble average conformational cost Δ*F*(Ξ), its entropy Δ*S*(Ξ), and then the free energy, according to Δ*G*(Ξ) = Δ*F*(Ξ) – *T*Δ*S*(Ξ) + Δ*G*_0_. The entropy Δ*S*(Ξ) is a measure for the diversity—the number of folded conformations that can be represented by the graph Ξ—of the subset. Δ*G*(Ξ) determines the overall thermodynamic stability of this subset relative to other subsets and the open chain. Δ*G*_0_ is the stabilization contributed by the duplexes in the structure. To determine the contribution from each duplex, we used the experimental Δ*G*_exp_ data reported by Sobzcak et al. for (CNG)_20_ oligomers in 100mM NaCl (50) for N = A, C, G and U as the target Δ*G*(Ξ). The conformation that was reported for (CNG)_20_ has the maximal hairpin structure shown in Fig. 4e—which is comprised of nine duplexes, eight symmetric (1,1) internal junctions, one 4-nt hairpin, and two dangling ends. The Δ*F*(Ξ) contributed by the loops of this structure was then calculated using our library of entropic costs: 5.10 kcal/mol for the 4-nt hairpin, 5.98 kcal/mol for each of the (1,1) internal junctions, and 0 for the dangling ends. As the maximal hairpin structure has no permutable segments on the junctions, there is one only possible structure that matches the graph in Fig. 4e. The stabilization contributed by each node is then calculated as 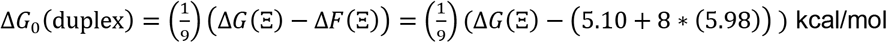. This calculation was carried out for each N = A, C, G and U using the experimental Δ*G*_exp_ data reported by Sobzcak et al. The smallest stabilization came from N = C with Δ*G*_0_(duplex) = −6.17 kcal/mol, followed by ∩ (Δ*G*_0_(duplex) = −6.39 kcal/mol), A (Δ*G*_0_(duplex) = −6.57 kcal/mol), and G (Δ*G*_0_(duplex) = −6.62 kcal/mol). In the following sections, we will use the N = C Δ*G*_0_(duplex) value to report our numerical data involving duplexes (black dots), as this represents a lower bound to stability for all the structures. The other values for N = A, G or T can be obtained by applying the appropriate offset to the values for each dot (duplex).

The graph ensemble calculations reported below have been carried out for a RNA strand total length *n* = 17 and 51. These were done by grouping graphs according to their total degree *D*, equal to the sum over the degrees of all vertices. Note that the number of graphs at each total degree *D* proliferates rapidly as *D* increases. However, the number of permissible graphs at each *D* is also constrained by the length of the RNA chain making it possible to exhaustively enumerate all graphs for chains that are not too long. In particular, for the *n* = 17 oligomer, we were able to enumerate all graphs up to *D* = 28, which is the highest-order permissible set. At *D* = 28, there is only one permissible graph, which is shown in Fig. 12(b), corresponding to the maximally base-paired structure for *n* = 17. All permissible graphs from *D* = 8 to 28 are displayed in Figs. 8 through 12 for the *n* = 17 oligomer (the *D* = 4 set is trivial and contains only one graph, which is not displayed). While exhaustively enumerating all graphs in the ensemble for *n* = 17 is possible, this quickly becomes impossible for longer repeat lengths. In the *n* = 51 case, the significantly longer chain length prevents the same total enumeration to be carried out, and only graphs up to total degree of *D* = 16 are presented in the data below.

**Figure 8.**
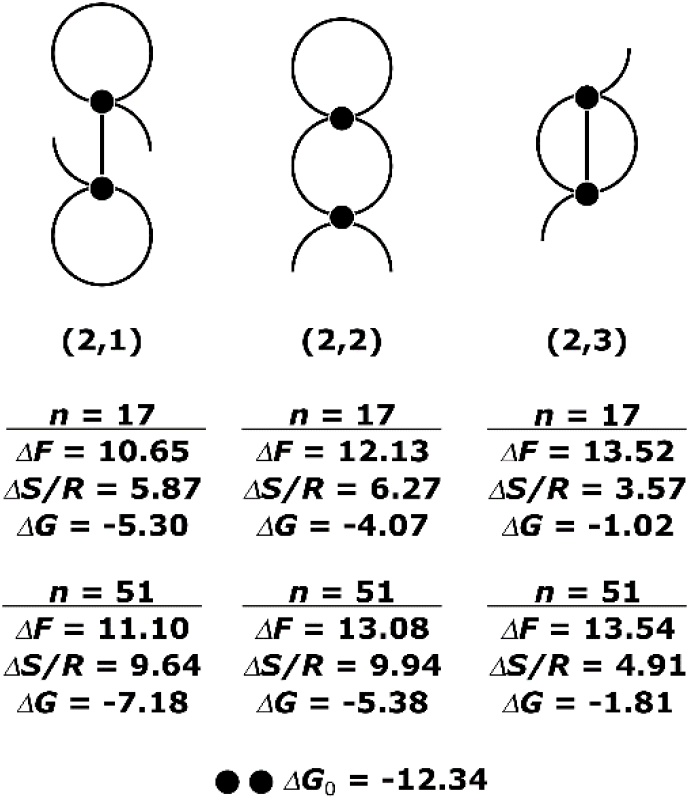
All graphs for (CNG)_17_ at total degree 8, their RAG-ID (19) and corresponding ensemble-averaged cost, entropy, and the graph free energy. Δ*F* and Δ*G* are in kcal/mol. Δ*S* are reported in units of *R*, the gas constant.

Fig. 8 shows all graphs with total degree *D* = 8 and the calculated values of Δ*F*(Ξ) and Δ*G*(Ξ) in kcal/mol, and Δ*S*(Ξ)/*R* for each. We derived these diagrams from the list of graphs enumerated by Schlick et al. (17, 19), after removing those containing structures for which we have no corresponding data or those requiring non-secondary structural motifs to form. Three motifs are present with total degree of 8: hairpins, two-way junctions, and pseudoknots. The thermodynamic stabilities of the three subsets reported in their Δ*G* values include the intrinsic stabilization provided by the free energy in the duplexes, Δ*G*_0_ = 2 × (−6.17) kcal/mol. Similarly, data are shown in Fig. 9 for all graphs with total degree *D* = 12 and in Fig. 10 for total degree *D* = 16. The stabilization provided by the duplexes are Δ*G*_0_ = 3 × (−6.17) kcal/mol and Δ*G*_0_ = 4 × (−6.17) kcal/mol, respectively. Figs. 11 and 12(a) show the non-pseudoknot graphs at *D* = 20,24 for *n* = 17, and Fig. 12(b) shows the single permissible graph at *D* = 28 which corresponds to a maximally base paired structure.

**Figure 9.**
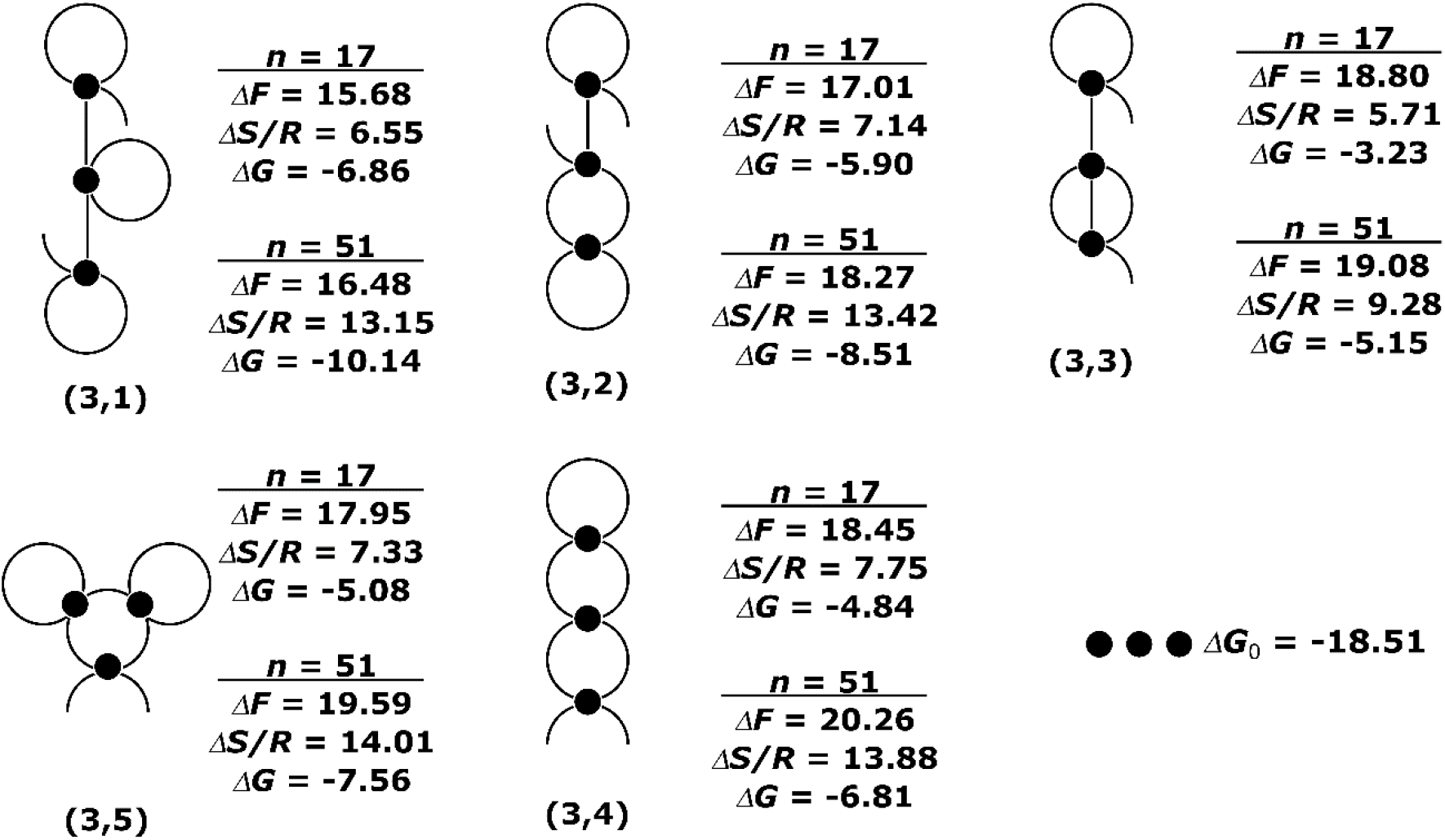
All graphs for (CNG)_17_ at total degree 12, their RAG-ID and corresponding ensemble-averaged cost, entropy, and the graph free energy. Δ*F* and Δ*G* are in kcal/mol. Δ*S* are reported in units of *R*, the gas constant.

**Figure 10.**
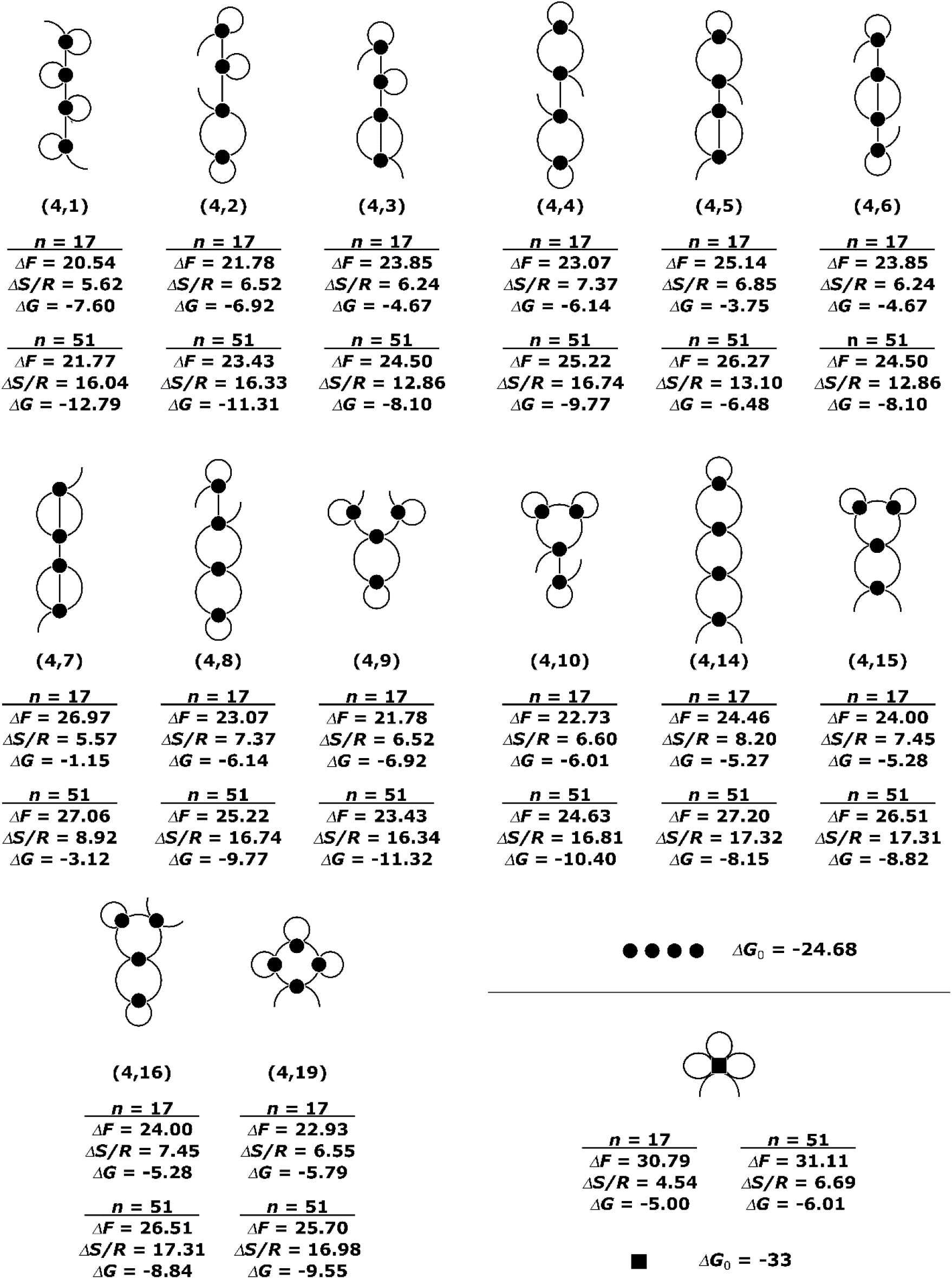
All graphs for (CNG)_17_ at total degree 16, their RAG-ID and corresponding ensemble-averaged cost, entropy, and the graph free energy. Δ*F* and Δ*G* are in kcal/mol. Δ*S* are reported in units of *R*, the gas constant. The list also includes a quadruplex structure, with an estimate for the intrinsic Δ*G* of the core, referenced against the same standard state (four 2-bp duplexes) used for the rest of the structures in this figure.

**Figure 11.**
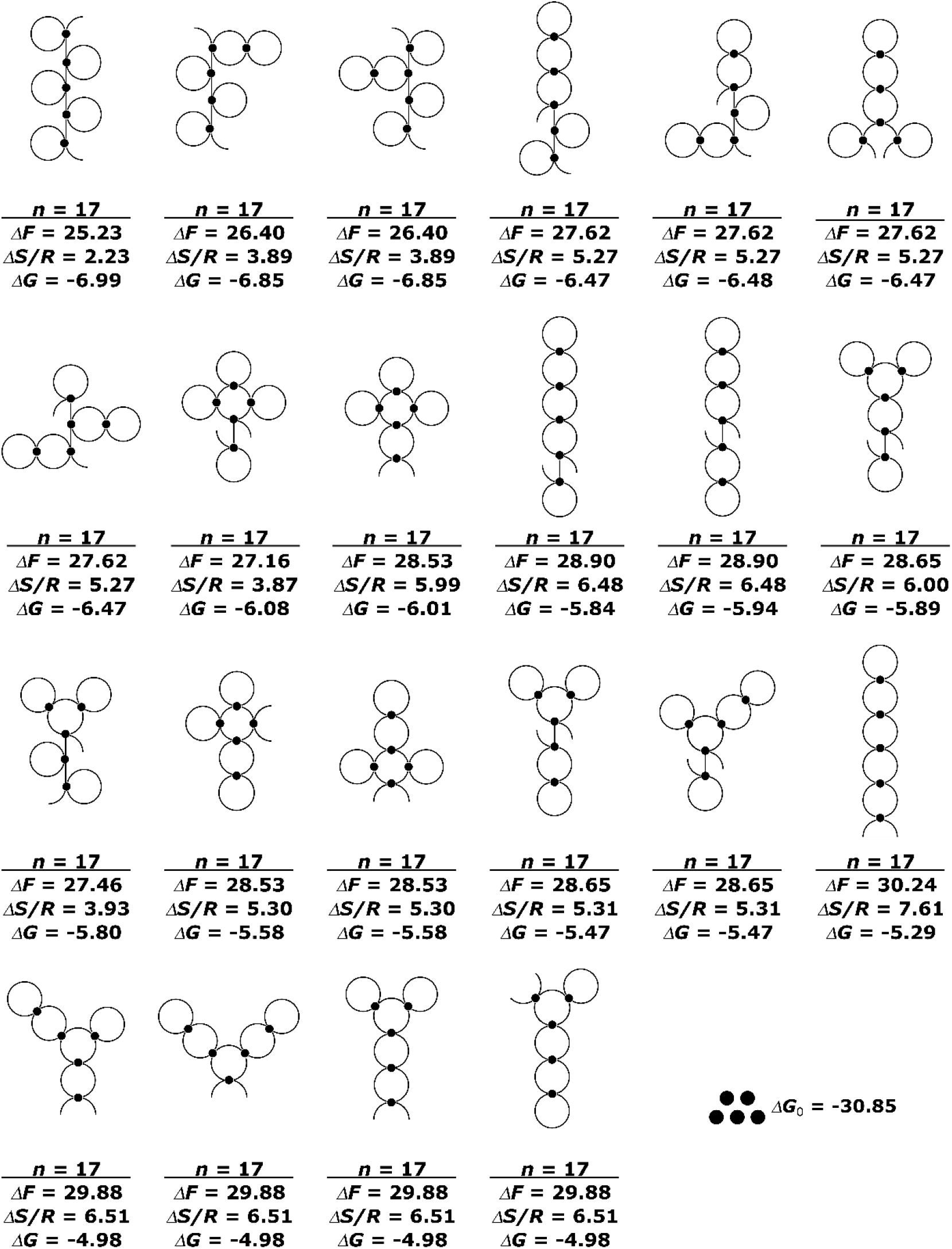
All non-pseudoknot graphs for (CNG)_17_ at total degree 20, their corresponding ensemble-averaged cost, entropy, and the graph free energy. Δ*F* and Δ*G* are in kcal/mol. Δ*S* are reported in units of *R*, the gas constant. The graphs have been sorted from most to least stable free energy rather than by RAG-ID

**Figure 12.**
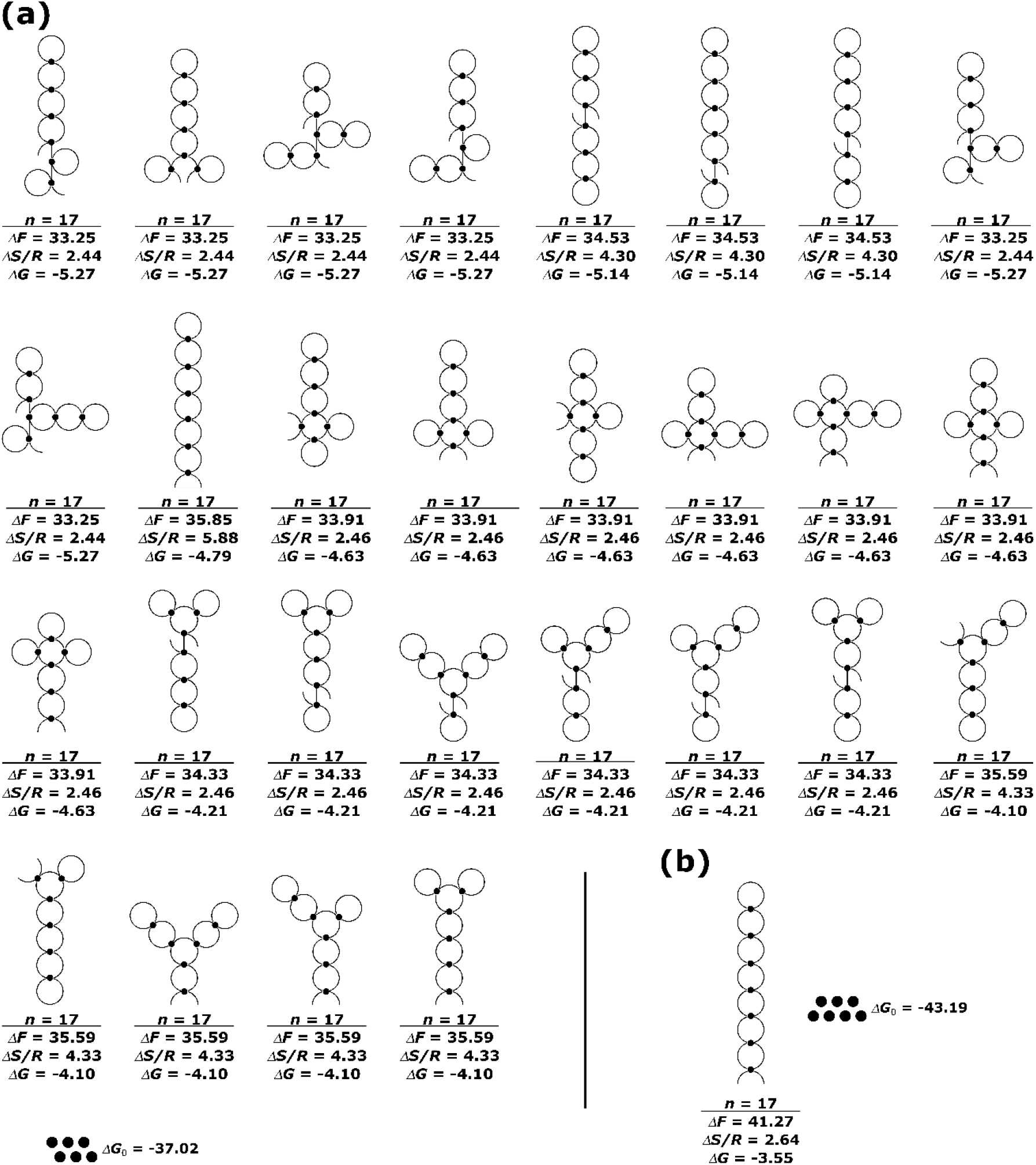
The single non-pseudoknot graphs for (CNG)_17_ at total degree 28 and all non-pseudoknot graphs of total degree 28, their corresponding ensemble-averaged cost, entropy, and the graph free energy. Δ*F* and Δ*G* are in kcal/mol. Δ*S* are reported in units of *R*, the gas constant. The graphs have sorted from most to least stable free energy rather than by RAG-ID. Note that due to the constraint of chain length, there are no bubble graphs with 6 or 7 nodes.

We also considered quadruplexes. The only graph that contains a single quadruplex is shown in Fig. 10. A single quadruplex has total degree *D* = 16. To establish the intrinsic free energy of the quadruplex core, we again rely on experimental data from Sobczak et al. Their study showed two trinucleotide oligomers which can form quadruplex in solution: (UGG)_17_ and (AGG)_17_, but (CNG) repeats cannot. To estimate the effects of including quadruplexes in the (CNG)_n_ repeat ensembles, we used the experimental free energies of (UGG)_17_ and (AGG)_17_. The proposed structure for both contains a single 2-layer quadruplex structure with a guanine tetrad in each layer similar to what is shown in Fig 6. The three loop lengths in the structure proposed by Sobczak et al. are all 1-nt in length. Like the maximal hairpin example above, we calculate Δ*G*(Ξ) for a quadruplex structure in a similar way, with only one single quadruplex node, which is represented in our graphs by a black square. Recall that Δ*G*(Ξ) = Δ*F*(Ξ) – *T*Δ*S*(Ξ) + Δ*G*_0_(quad), and the loop contribution in the form of Δ*F*(Ξ) is comprised of the cost for forming each of the three linker loops a, b, and c, and the cost of aligning the guanine in each of the tetrad’s column. As only one quadruplex structure is reported in the work of Sobczak et al—a quadruplex on the 5’ end and a long terminal tail—Δ*S*(Ξ) is zero. The cost of loop *a*, loop *b*, and aligning all columns in the quadruplex core are read directly from Table 1: 4.2 kcal/mol, 5.3 kcal/mol, and 12.6 kcal/mol, respectively. The contribution of loop *c* is approximated at 7.0 kcal/mol due to lack of explicit simulated value at 1-nt. The free energy of a quadruplex core, Δ*G*_0_(quad), is then calculated as Δ*G*_0_(quad) = Δ*G*(Ξ) – Δ*F*(Ξ) = Δ*G*(Ξ) – (4.2 + 5.3 + 7.0 + 12.6) kcal/mol. Experimentally determined Δ*G*_exp_ = Δ*G*(Ξ) for (UGG)_17_ and (AGG)_17_ in 100mM NaCl at 260nm from Sobzcak et al. were used, yielding an approximation for the quadruplex core of −32.97 kcal/mol from (AGG)_17_ and −33.14 kcal/mol from (UGG)_17_. The value −33 kcal/mol was used in calculating the free energy of the single quadruplex-containing graph in Fig. 10.

## DISCUSSION

Data in Figs. 8–12 reveal the basic characteristics of the structural ensembles typical of CNG repeat sequences. While this direct enumeration approach is limited to graphs of low total degrees and/or relatively short total chain lengths, the results demonstrate central features that allow us to make projections about graphs of higher degrees and longer total chain length. We begin with a discussion of the graph ensembles for 5’-NG(CNG)_16_CN-3’ (*n* = 17).

At total degree *D* = 8, Fig. 8 reveals that the graph with the lowest overall free energy is (2,1). While this is just one graph, it is important to remember that a large number of conformations are represented by it, where the segments in the graph can have variable lengths but they are restricted in such a way that their sum must equal the total length of the full RNA chain. The entropy Δ*S/R* listed under the graph is a measure for the number of these conformations. The value Δ*S/R* = 5.87 for graph (2,1) when *n* = 17 suggests that there are roughly *e*^5.87^~354 conformations represented. The value Δ*F* is the sub-ensemble free energy cost associated with suppressing the backbone conformational degrees of freedom to force the chain to conform with the constraints implied by the vertices in the graph. For (2,1) it is 10.65 kcal/mol when *n* = 17. Notice that for every conformation in this sub-ensemble the backbone conformational cost is different, and the conformation tally of ~354 as well as the cost Δ*F* = 10.65 kcal/mol are ensemble averaged properties. The overall free energy of graph (2,1) when *n* = 17 is Δ*F* – (*RT*)(ΔS/R) + ΔG_0_ = 10.65 – (0.616)(5.87) + Δ*G*_0_ = −5.30 kcal/mol, where Δ*G*_0_ is the intrinsic free energy associated with two 2-bp duplexes, equal to −12.34 kcal/mol as indicated in Fig. 8, and *T* = 310 K.

The graph that has the next highest overall free energy in Fig. 8 is (2,2), with Δ*G* = −4.07 kcal/mol. The entropy of graph (2,2) is similar to (2,1). This is because their topologies are similar, except two segments in (2,2) are constrained into a 2-way junction, whereas in (2,1) one of them is constrained inside a hairpin and the other is free. Contrasting this to the graph (2,3) in Fig. 8, which contains a pseudoknot, Δ*S/R* is quite a bit lower for (2,3), even though (2,3) contains the same number of segments as (2,2). Note that Eq. (1) dictates that the total number of edges *E* and the total degree *D* are related by *E* = 1 + *D*/2, and all graphs in Fig. 8 necessarily have the same number of edges. The reason why the structure containing the pseudoknot has a significantly lower entropy compared to (2,1) and (2,2) is related to the dependence of the cost function of a pseudoknot on the length of the unpaired loop regions that comprise it. Long loop lengths are suppressed in a pseudoknot compared to hairpins or 2-way junctions, and this leads to a lower diversity in the sub-ensemble associated with graph (2,3) compared to (2,1) or (2,2). This also results in a higher overall Δ*G* for graphs containing pseudoknots.

While each graph represents a sub-ensemble of the conformations at a certain total degree *D* and its entropy value Δ*S/R* reflects the diversity of that subset, an additional ensemble-level entropy is associated with the superset of graphs at each *D*. This ensemble-level entropy at a degree *D* is given by Δ*S_D_*/*R* = − Σ_Ξ_*P*(Ξ) ln *P*(Ξ), where the normalized probability *P*(Ξ) of graph Ξ is given by 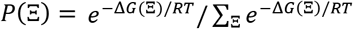. This value can be interpreted as the measure of how all the conformation at the *D* = 8 level is distributed amongst the graphs in Fig. 8. For the graphs in Fig. 8 with *n* = 17, Δ*S_D_*/*R* comes out to be 0.37, suggesting that at the ensemble level, the information content in the superset of *D* = 8 graphs is roughly equivalent to just *e*^0.37^~1.4 graphs. This suggests that the secondary structural content attributed to the graphs in Fig. 8 are not evenly distributed amongst the three graphs. As Δ*G*(Ξ) determines the weight of each graph, this suggests that most of the configurations are consistent with the non-pseudoknot graphs. Finally, the overall free energy of the ensemble Δ*G_D_* can be computed either from 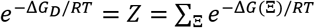 or Δ*G_D_* = 〈Δ*G*〉 – *T*Δ*S_D_*, where 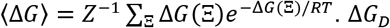 serves as a measure for thermodynamic stability of the *D* = 8 sub-ensemble—that is, the collection of all *D* = 8 structures—relative to the open chain. For the *D* = 8 sub-ensemble, Δ*G_D_* = −5.38 kcal/mol. This suggests that the *D* = 8 graphs are collectively stable relative to an open chain.

Moving to the *D* = 12 graphs in Fig. 9, the diversity of the graphs expands. The member of this superset with the lowest overall free energy Δ*G* is (3,1). The one with the highest free energy is again the structure with a pseudoknot. *D* = 12 is the lowest order at which a 3-way junction appears such as in (3,5). Similar to the *D* = 8 superset in Fig. 8, the member with the highest sub-ensemble diversity is (3,4), which has the highest entropy Δ*S/R* = 7.75, corresponding to approximately *e*^7.75^~2300 distinct conformations. The ensemble-level entropy for the set *D* = 12 is Δ*S_D_/R* = 0.75 corresponding to ~ 2.1 graphs. The overall free energy of the *D* = 12 sub-ensemble is Δ*G_D_* = −7.02 kcal/mol. The values show that there are more graphs that comprise the *D* = 12 sub-ensemble and collectively they are more favorable than the *D* = 8 sub-ensemble.

Going to *D* = 16 in Fig. 10, the diversity of the graphs expands even further. All the graphs in Fig. 10 contain four 2-bp duplexes, except the one with a quadruplex. The intrinsic free energy associated with four 2-bp duplexes is Δ*G*_0_ = −24.68 kcal/mol, which is indicated in Fig. 10. On the other hand, the intrinsic free energy of a 2-layer quadruplex core, according to the reasoning at the end of the last section, is estimated to be Δ*G*_0_~−32.97 kcal/mol. Fig. 10 shows that the Δ*G* value for the quadruplex graph is −4.97 kcal/mol. Its entropy is low because of a reason similar to the pseudoknots – the quadruplex structure is confined to short loop lengths and reduces the conformational diversity of the subset. Notice that the quadruplex structure is only possible when the sequence is (CGG)_M_ and thus is not expected to contribute significantly to the structural ensemble of (CNG) repeats.

Like the graphs in Figs. 8 and 9, the graph with the lowest overall free energy in Fig. 10 is graph (4,1). We will refer to graphs having this topology as “bubble diagrams”. For graph (4,1), the overall free energy is Δ*G* = −7.60 kcal/mol. Notice that going from *D* = 12 in Fig. 9 to *D* = 16 in Fig. 10, the entropy of the bubble diagram actually decreases from Δ*S/R* = 6.55 for graph (3,1) in Fig. 9 to 5.62 for graph (4,1) in Fig. 10. This indicates that when going to higher total degree *D*, the fixed length of the full RNA chain becomes a factor limiting the number of combinations of segment lengths that could fit into the total number of nucleotides on the sequence. However, this effect is also dependent on other features of the graphs. For example, the type of graphs with the highest entropy in both Fig. 9 and 10 are the “necklace diagrams”, exemplified by structures (3,4) and (4,14). Going from (3,4) to (4,14), the entropy value of the necklace diagram continues to increase from Δ*S/R* = 7.75 to 8.20, growing from ~2300 to ~3600 configurations. Furthermore, some of the other partial necklace diagrams, such as (3,2) in Fig. 9 and (4,4) and (4,8), also show continued increase in diversity going from lower to higher order total degree. In addition to these, the additional diagrams associated with 3- or 4-way junctions also seem to expand in diversity. The ensemble-level entropy for the set *D* = 16 is Δ*S_D_/R* = 1.64 corresponding to ~ 5.2 graphs, and comparing this to Δ*S_D_/R* =0.67 for *D* = 12, validates this observation. The overall free energy of the *D* = 16 sub-ensemble is Δ*G_D_* = −8.05 kcal/mol, indicating that the superset of *D* = 16 diagrams continues to be more thermodynamically favorable than the graphs of lower total degree.

Going beyond *D* = 16, Fig. 11 shows all non-pseudoknot graphs at *D* = 20 containing up to 4-way junctions. Fig. 12a shows all non-pseudoknot graphs at *D* = 24. Fig. 12b shows the single nonpseudoknot graph at *D* = 28. For *D* > 16, pseudoknot graphs have been omitted due to their falloff in thermodynamic stability and the small sub-ensembles they represent. Our graphs also do not include multiway junctions higher than 4 because our library does not contain simulation data for 5-way or higher junctions. For *n* = 17 chains, however, only one possible graph at *D* = 28 would have a 5-way junction. For *n* = 17 chains, there are no graphs beyond *D* = 28.

In the *D* = 20 graphs in Fig. 11, the bubble diagram continues to be the most energetically favorable. However, the *D* = 20 graphs collectively are no more favorable than the graphs of lower degrees. Most of them have ΔG similar to graphs in Fig. 10, and some of them are less favorable. The *D* = 20 subensemble entropy is *S_D_/R* = 2.66, corresponding to ~14.4 graphs, and the free energy is Δ*G_D_* = −8.10. All of the non-pseudoknot graphs in Fig. 10 at *D* = 16 have four duplexes (dots), whereas the graphs in Fig. 11 at *D* = 20 have five. Our results show that relative to the open chain, 5-duplex structures collectively are only marginally more favorable than 4-duplex structures. This trend continues in Fig. 12a for *D* = 24 where all the graphs are now less thermodynamically stable than their counterparts at lower degrees. This decrease in thermodynamic stability is an intrinsic entropic effect produced by the total length of the RNA chain which constrains the number of permutable loop segments. As more nodes are added for a given total chain length, fewer free nucleotides are available to be assigned to the to the unpaired loop regions. This leads to a decrease in the structural diversity Δ*S/R* that offsets the entropic cost of constraining the backbone. This can be seen in the Δ*S/R* values of the graphs going from *D* = 16 to *D* = 20 and beyond. The *D* = 24 sub-ensemble has an entropy Δ*S_D_/R* = 3.01, corresponding to ~20.3 graphs and free energy Δ*G_D_* = −6.84. These values indicate that despite more graphs of total degree *D* = 24 being present, the 6-node structure corresponding to them are less favorable collectively than the 5-node and 4-node graphs. The chain length further constrains the graph down to a single diagram in Fig. 12b for *D* = 28, which is the maximum degree possible for *n* = 17. This diagram has only one permutable segment and hence a low entropy.

Data for the *n* = 51 chain demonstrate the effects chain length exerts on the characteristics of the graph sub-ensembles. Figs. 8–10 show that the average cost ΔF of all graphs increases when the chain grows from *n* = 17 to *n* = 51. This is expected as longer RNA chains should have access to secondary structures with longer loop lengths. However, the changes in cost reflect a change in loop length of only 3 to 6 nucleotides, and the secondary structures of the graphs only grew by one or two additional trinucleotide units as the total RNA chain length is tripled. This points to a preferential placement of nucleotide units into the dangling ends or bridging junctions which incur no cost of formation while contributing to the overall structural diversity of the graph.

At the same time, the increase in RNA chain length has a large favorable effect on the graphs’ entropies. Every graph exhibited an increase in Δ*S/R* when the chain length grew from *n* = 17 to 51. This is also expected as there are now more transposable units and the RNA chain has access to a larger number of conformations for each graph. The changes in Δ*F* and Δ*S* combine to yield more favorable Δ*G* for all graphs as the total RNA chain length grows. The longer total RNA chain length also increases the maximal number of nodes that can be present in the graphs. Though we are not able to investigate the graphs of total degree greater than 16 for the *n* = 51 chain, the results from the *n* = 17 case suggests that at the Δ*G* of the graphs will peak at some total degree near the maximum, which for *n* = 51 is *D* = 96 or 24 nodes.

Our data suggest that as a function of repeat length *n*, there are two opposing factors that control the thermodynamic stability of the graphs at different degrees *D*. First, longer repeat lengths permits a larger number of duplexes to be made, and the maximum degree *D^‡^* is proportional to *n*. The duplexes contain extra thermodynamic stability due to the base pairs and base stacks, and graphs at higher *D* have more node stabilization. But at the same time, more duplexes constrain the backbone conformations, producing lower conformational entropy for the chains. This acts against larger *D* and destabilizes them. Finally, the repeat length *n* also limits the number of possible diagrams as *D* increases toward the maximum *D^‡^*. This further truncates the size of the sub-ensembles approaching *D^‡^*. This tradeoff between duplex stability and chain conformational diversity results in an optimum in the sub-ensemble free energy Δ*G_D_* at a certain *D*. Stronger intrinsic duplex stability shifts this optimum position toward *D^‡^*, whereas a weaker duplex stability shifts this optimum position toward lower *D*. Depending on the width of this distribution, the ensemble may be characterized by graphs from many different degrees *D* with open diagrams being the most common dominant sub-ensemble.

These theoretical predictions are in variance from the prevailing view that the dominant structure of CNG repeats is a maximal hairpin structure with 2-way junctions plus a single hairpin. The results instead point to many potential structures of similar prevalence with large contribution from open and bubble-diagram type structures. Though crystallographic data point to the dominance of hairpin structures, it leaves the question of how an ensemble of mostly open structures can be detected in solution. Techniques such as small-angle X-ray scattering (SAXS) (51–54), UV melting (55), and Forster resonance energy transfer (FRET) (56) can all be used to probe the solution structure of RNA. While the use of thermodynamic data of Sobzcak et al. does provide a point of contact between the calculated free energies and experimental measurements, the ensemble predicted by our results is diverse enough that a one-to-one correspondence to experiments is unlikely. Instead, general features consistent with a diverse ensemble of open and flexible structure should be looked for in the experimental data to confirm our prediction. In particular, the predicted ensemble should have an SAXS profile yielding Kratky plot consistent with flexible structures, UV melting data consistent with a broad structural distribution, and FRET measurements that show contacts between positions proximal to one another with stretches of no contacts in between.

We now address possible limitations of our model in its current form and future steps to improve on what we have presented here. As the focus of this study was to understand the diversity of (CNG) repeats at a secondary structural level, long range interactions were not included. These studies can be carried out using the same Nucleic MC simulation framework to measure the free energy of these tertiary contacts, and these are currently in progress. Additionally, the current model assumes WC-only base pairing, and as such, ignores base pairs between the non-GC residues in the repeats. However, the enumeration of diagrams involving non-WC pairs is possible, and a library of free energies of these contacts can be constructed using the same MC simulation framework to compute their free energies, and these are also currently underway. Also enumerating diagrams with non-WC pairs introduce additional graph elements which also makes the enumeration process more complex, but this limitation can be addressed by the diagrammatic summation method proposed in the companion paper, wherein we apply a partition function method to study the graph ensembles, removing the need for explicit enumeration (57).

## CONCLUSION

Our analysis of the graph ensembles of CNG repeat chains at the oligomer scale has provided a first look at a theoretical model for analyzing the structural diversity of trinucleotide repeat chains, as well as observations that will be germane to understanding RNA conformational ensemble. By using a graph factorization method and a data library that has been built from simulations, we were able to group accessible secondary structures together into subsets represented by graphs and have calculated metrics for their thermodynamic stability Δ*G*(Ξ), as well as the structure content Δ*S*(Ξ) of the sub-ensembles. The results show that most structures are thermodynamically stable with a range of stability which would result in the prevalence of certain structures over others. The addition of helices—corresponding in our data to an increase in the total degree of a graph—incurs an additional cost that is offset by an increase in the number of structure accessible to the chain, and it is the balance between these two factors that determines the thermodynamic favorability of the structure. When the total degree (helices) increases, the dominant conformations are associated with the so-called bubble diagram—graphs composed of only hairpins connected by bridging unpaired segments, while the necklace diagrams tend to have higher entropy but also higher free energy costs. The results show that the extent to which the structural diversity of different classes of diagrams can grow as the total degree increases is also dictated by the chain length and the stabilization provided by the helices present in the structure. Some structures, such as the bubble diagrams, begin to lose structural diversity as the total degree grows past *D* = 16 for short chains, while others continue to proliferate.

Altogether, the results show that the structural diversity and propensities for different structural elements on CNG repeat chains are determined by an interplay between the length of the full RNA chain, the stabilizing strength of the helices, and the complexity of the graphs in the ensemble, In future studies, we will continue to explore different facets of these questions to understand the structural diversity and stability of long trinucleotide repeat chains on the length scale associated with the onset of TREDs.

## AUTHOR CONTRIBUTIONS

CHM designed the study. ENHP carried out simulations, data collection and the calculations. ENHP and CHM wrote the manuscript.

## ACKNOWLEDGEMENTS

This material is based in part upon work supported by the National Science Foundation under Grant Number CHE-1664801.

The authors also thank Peter Z. Qin for helpful discussions and comments.

## Notes

### Competing Interest Statement

The authors have declared no competing interest.

### Summary of Updates

The Results section has been expanded to include ensemble of chains up to (CNG)_29_ and previously presented ensembles have been fully enumerated. New results for (CNG)_51_ were added to approximate the critical expansion threshold observed in TRED. Additionally thermodynamic values used in the model have been recalibrated with respect to experimental data to allow more direct contact between the calculations using the model and with experimental values.

