## Supplemental information for "Quantifying Structural Diversity of CNG Trinucleotide Repeats Using Diagrammatic Algorithms"

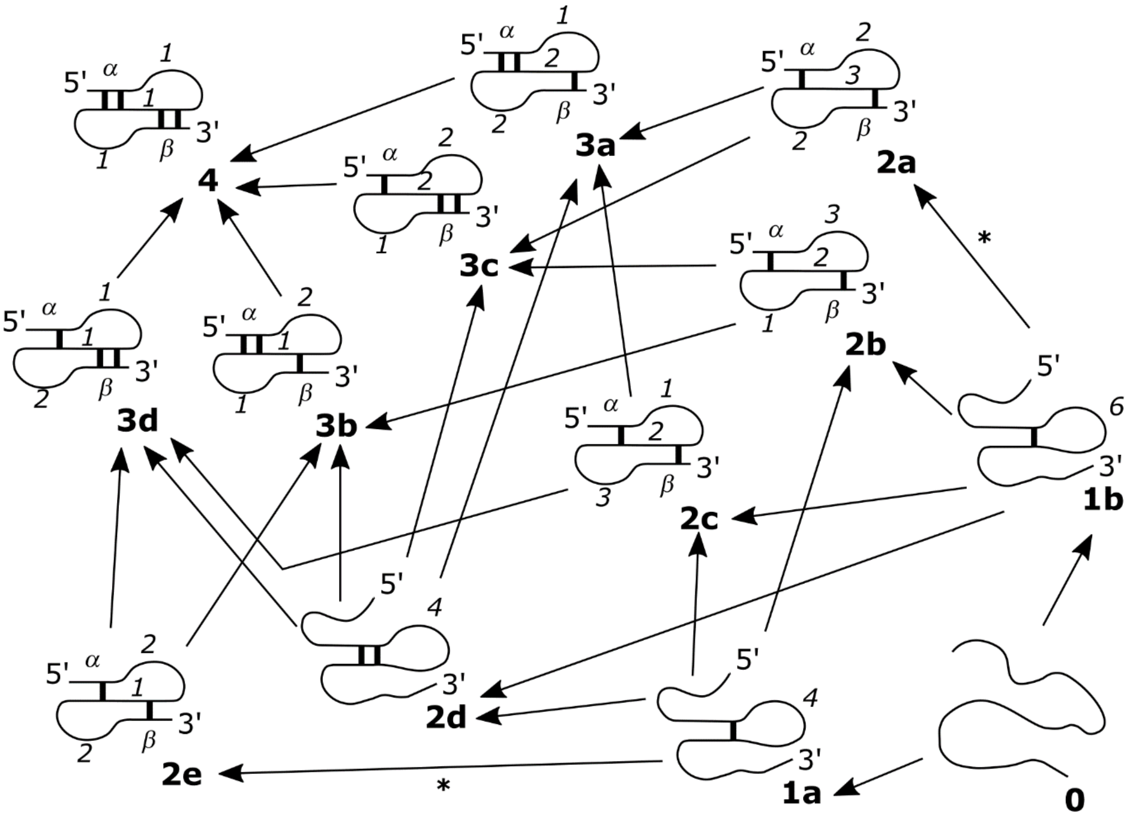


#### Figure S1.

Map showing the thermodynamic pathways used in calculating the entropic cost to form a pseudoknot structure with the junction length *a=b=c=1*. The entropic cost associated with each step can be found in Table S1. Steps leading to symmetric structures in which the choice of 5' or 3' for the base pairing event is indistinguishable are marked with an asterisk and have entropic cost reported for both the 5' and 3' base pairing.

| Start | End | Cost | Start | End | Cost | Start | End | Cost |
|---|---|---|---|---|---|---|---|---|
| 0 | 1a | 5.01 | 2a | 3a | 6.03* | 3a | 4 | 6.03* |
| 0 | 1b | 5.61 | 2a | 3c | 6.03* | 3b | 4 | 6.75* |
| 1a | 2b | 5.82 | 2b | 3b | 6.03* | 3c | 4 | 6.03* |
| 1a | 2c | 6.00 | 2b | 3c | 6.75* | 3d | 4 | 6.75* |
| 1a | 2d | 5.57 | 2c | 3a | 6.75* | | | |
| 1a | 2e (5') | 5.11 | 2c | 3d | 6.03* | | | |
| 1a | 2e (3') | 5.21 | 2d | 3a | 7.46 | | | |
| 1b | 2a (5') | 5.77 | 2d | 3b | 6.34 | | | |
| 1b | 2a (3') | 5.82 | 2d | 3c | 7.46 | | | |
| 1b | 2b | 5.26 | 2d | 3d | 6.38 | | | |
| 1b | 2c | 5.22 | 2e | 3b | 6.75* | | | |
| 1b | 3d | 4.85 | 2e | 3d | 6.75* | | | |

**Table S1. Entropic Cost for the Stepwise Formation of the *a=b=c=1* Pseudoknot**

Cost of each step in the thermodynamic pathways of Figure S1. Numbers marked with an asterisk are obtained via closed path calculations rather than directly obtained from simulations. All costs are reported in units of kcal/mol

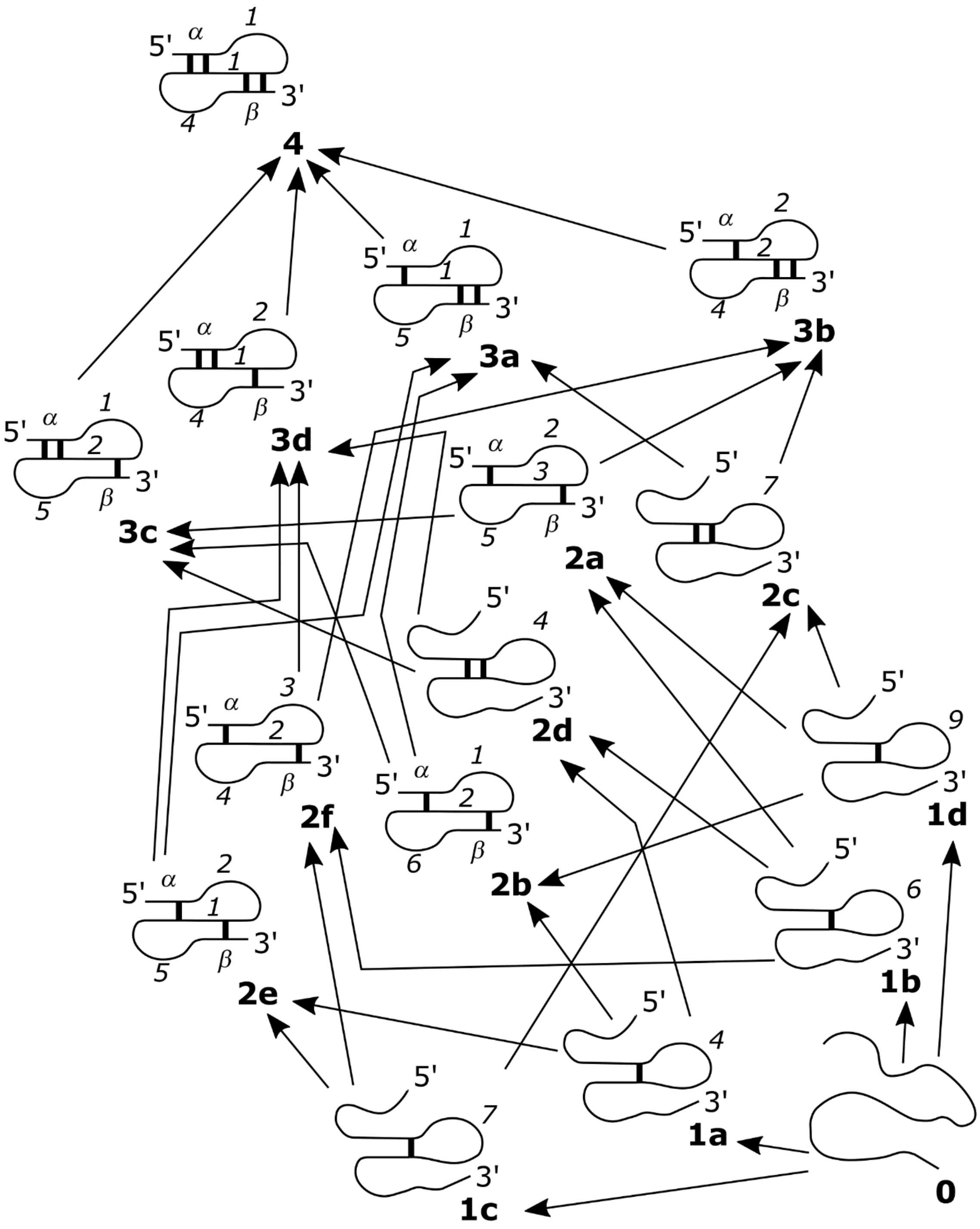


**Figure S2.**

Map showing the thermodynamic pathways used in calculating the entropic cost to form a pseudoknot structure with the junction length *a=b=1, c=4*. The entropic cost associated with each step can be found in Table S2.

| Start | End | Cost | Start | End | Cost |
|---|---|---|---|---|---|
| 0 | 1a | 5.02 | 2a | 3b | 6.03* |
| 0 | 1b | 5.62 | 2a | 3c | 6.03* |
| 0 | 1c | 5.85 | 2b | 3a | 6.03* |
| 0 | 1d | 6.20 | 2b | 3c | 6.75* |
| 1a | 2b | 6.39 | 2c | 3a | - |
| 1a | 2d | 5.51 | 2c | 3b | - |
| 1a | 2e | 5.94 | 2d | 3c | 7.71 |
| 1b | 2a | 6.63 | 2d | 3d | 7.37 |
| 1b | 2d | 4.85 | 2e | 3a | 6.75* |
| 1b | 2f | 6.18 | 2e | 3d | 6.75* |
| 1c | 2c | 5.51 | 2f | 3b | 6.75* |
| 1c | 2e | 5.15 | 2f | 3d | 6.03* |
| 1c | 2f | 5.86 | 3a | 4 | 6.75* |
| 1d | 2a | 6.05* | 3b | 4 | 6.03* |
| 1d | 2b | 5.21* | 3c | 4 | 6.03* |
| 1d | 2c | 4.85 | 3d | 4 | 6.75* |

**Table S2. Entropic Cost for the Stepwise Formation of the *a*=*b*=*1, c*=*4* Pseudoknot**

Cost of each step in the thermodynamic pathways of Figure S2. Numbers marked with an asterisk are obtained via closed path calculations rather than directly obtained from simulations. Steps for which no simulation data nor sufficient data for closed-path calculations were available have been left blank. All costs are reported in units of kcal/mol

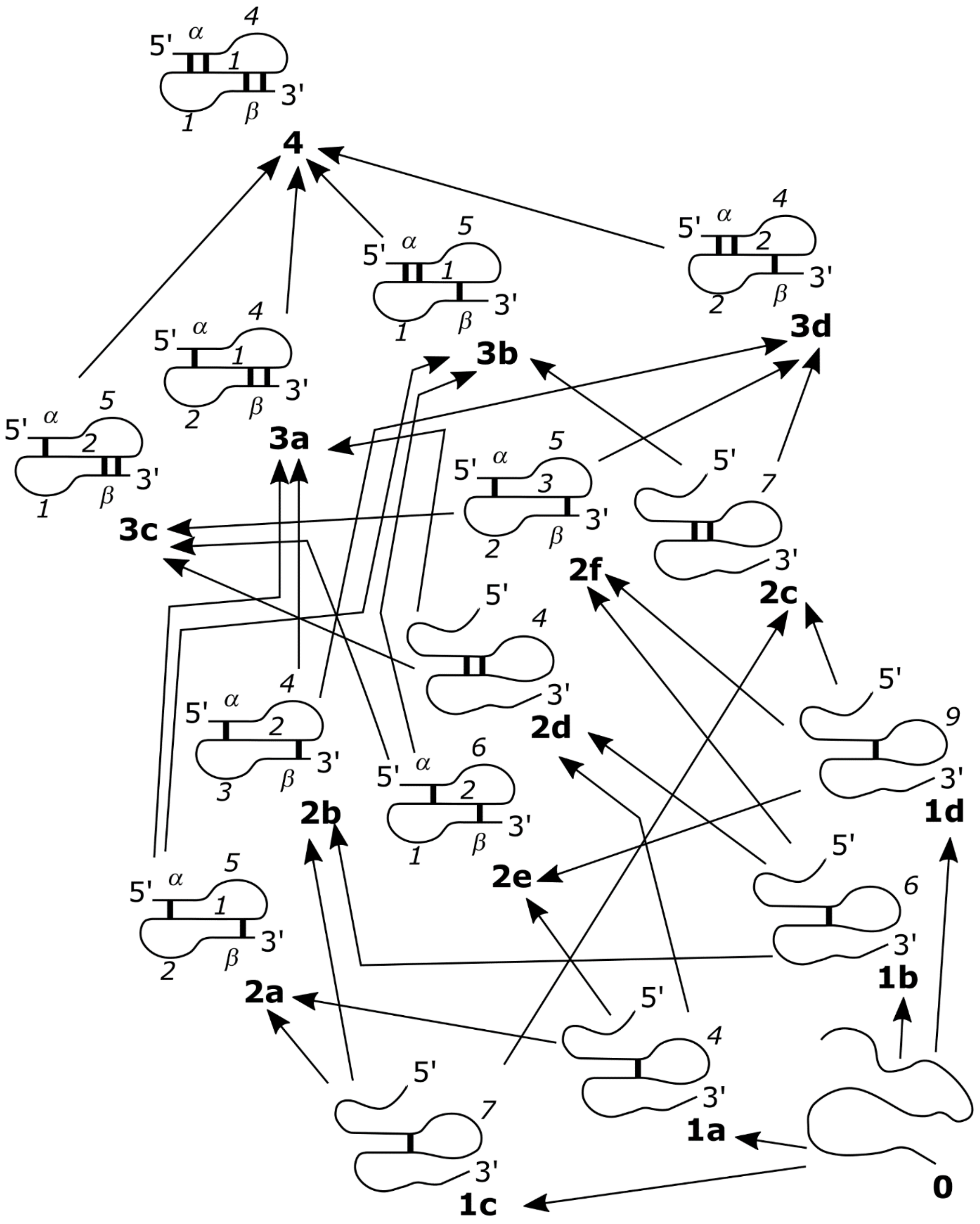


**Figure S3.**

Map showing the thermodynamic pathways used in calculating the entropic cost to form a pseudoknot structure with the junction length *a=4, b=c=1*. The entropic cost associated with each step can be found in Table S3.

| start | end | cost | start | end | cost |
|---|---|---|---|---|---|
| 0 | 1a | 5.02 | 2a | 3a | 6.75* |
| 0 | 1b | 5.62 | 2a | 3b | 6.75* |
| 0 | 1c | 5.85 | 2b | 3a | 6.03* |
| 0 | 1d | 6.20 | 2b | 3d | 6.75* |
| 1a | 2e | 6.39 | 2c | 3a | - |
| 1a | 2a | 5.98 | 2c | 3b | - |
| 1a | 2d | 5.51 | 2d | 3a | 6.86 |
| 1b | 2b | 6.08 | 2d | 3c | 8.14 |
| 1b | 2d | 4.85 | 2e | 3b | 6.03* |
| 1b | 2f | 6.51 | 2e | 3c | 6.03* |
| 1c | 2a | 5.20 | 2f | 3c | 6.75* |
| 1c | 2b | 5.90 | 2f | 3d | 6.03* |
| 1c | 2f | 5.51 | 3a | 4 | 6.75* |
| 1d | 2a | - | 3b | 4 | 6.75* |
| 1d | 2b | - | 3c | 4 | 6.03* |
| 1d | 2c | - | 3d | 4 | 6.03* |

#### Table S3. Entropic Cost for the Stepwise Formation of the *a=4, b=c=1* Pseudoknot

Cost of each step in the thermodynamic pathways of Figure S2. Numbers marked with an asterisk are obtained via closed path calculations rather than directly obtained from simulations. Steps for which no simulation data nor sufficient data for closed-path calculations were available have been left blank. All costs are reported in units of kcal/mol

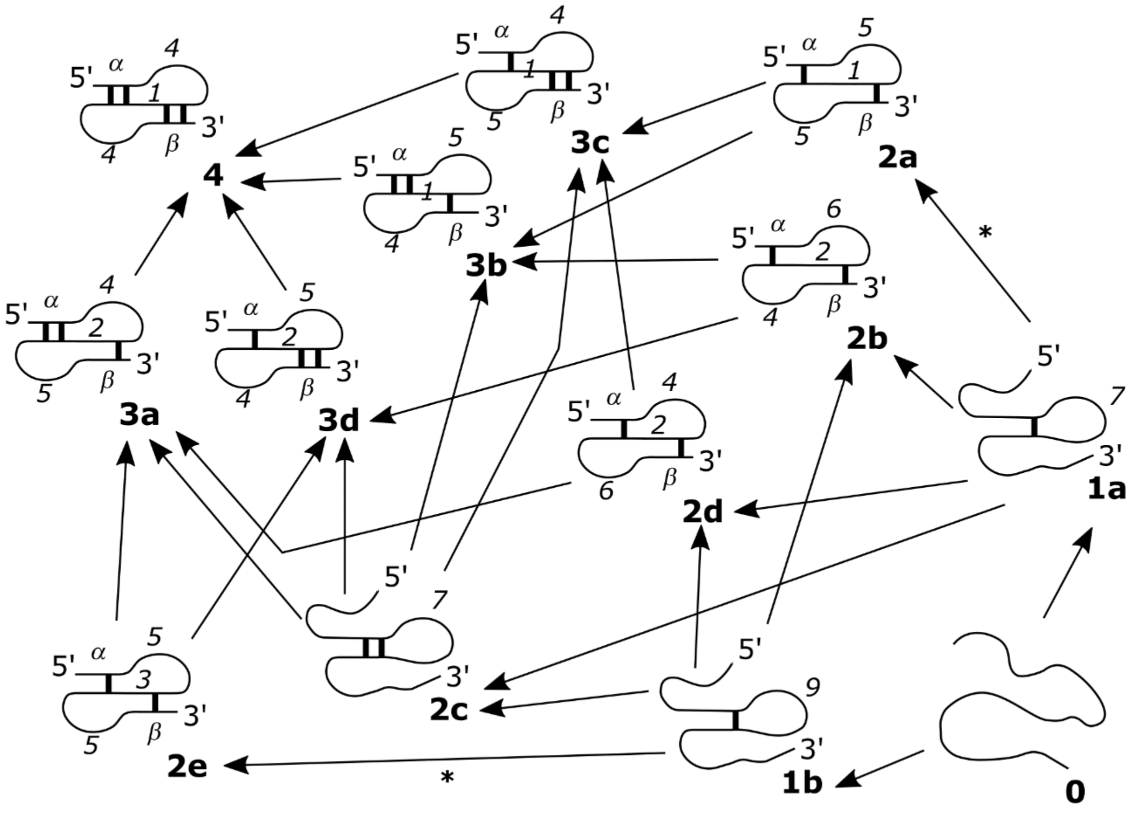


**Figure S4.**

Map showing the thermodynamic pathways used in calculating the entropic cost to form a pseudoknot structure with the junction length *a=c=4, b=1*. The entropic cost associated with each step can be found in Table S4. Steps leading to symmetric structures in which the choice of 5’ or 3’ for the base pairing event is indistinguishable are marked with an asterisk and have entropic cost reported for both the 5’ and 3’ base pairing. Note that structure 1b and 2c do not have many available values and were omitted from the pathway analysis. Consequently 2e is also not included due to being accessible only from 1b. These structures have been included here for the sake of completion.

| start | end | cost | start | end | cost |
|---|---|---|---|---|---|
| 0 | 1a | 5.85 | 2a | 3b | 6.75* |
| 0 | 1b | 6.20 | 2a | 3c | 6.75* |
| 1a | 2a (5') | 5.97 | 2b | 3b | 6.03* |
| 1a | 2a (3') | 5.90 | 2b | 3d | 6.75* |
| 1a | 2b | 6.38 | 2c | 3a | - |
| 1a | 2c | - | 2c | 3b | - |
| 1a | 2d | 6.26 | 2c | 3c | - |
| 1b | 2b | - | 2c | 3d | - |
| 1b | 2c | - | 2d | 3a | 6.75* |
| 1b | 2d | - | 2d | 3c | 6.03* |
| 1b | 2e | - | 3a | 4 | 6.03* |
| | | | 3b | 4 | 6.75* |
| | | | 3c | 4 | 6.75* |
| | | | 3d | 4 | 6.03* |

### Table S4. Entropic Cost for the Stepwise Formation of the *a=c=4, b=1* Pseudoknot

Cost of each step in the thermodynamic pathways of Figure S2. Numbers marked with an asterisk are obtained via closed path calculations rather than directly obtained from simulations. Steps for which no simulation data nor sufficient data for closed-path calculations were available have been left blank. Structure 2e has been fully omitted from the table as no simulation data for its formation from a prior structure nor the subsequent steps leading out from it is available.
